# In vivo imaging of inferior olive neurons reveals roles of co-activation and cerebellar feedback in olivocerebellar signaling

**DOI:** 10.1101/2025.01.06.631443

**Authors:** Da Guo, Marylka Yoe Uusisaari

## Abstract

Complex spikes (CSs), generated by inferior olive (IO) neurons, are foundational to most theories of cerebellar function and motor learning. Despite their importance, recordings from IO neurons in living animals have been limited to single-electrode methods, providing no insights into multineuron dynamics within intact circuits. Here, we used a novel ventral surgical approach that allows calcium imaging-based monitoring of multicellular activity in the IO of anesthetized mice. This method provides direct optical access to the ventral medulla, enabling simultaneous recording of spontaneous and sensory-evoked activity within localized clusters of IO neurons, specifically in the principal (PO) and dorsal accessory olives (DAO).

Our findings reveal that spontaneous activity rates and event magnitudes differ between the PO and DAO, consistent with observations from cerebellar cortex recordings in zebrin-positive (Z+) and zebrin-negative (Z-) zones, respectively. We further demonstrate that spontaneous event amplitudes are influenced by co-activation among neighboring neurons, so that events occurring in clusters are larger than single ones. Event co-activation is more pronounced in the PO than in the DAO, potentially explaining the differences in complex spike sizes observed in Z+ and Z-zones. Sensory-evoked events induced by periocular airpuff stimulation were larger than spontaneous ones, as expected. However, this difference diminishes when accounting for the higher levels of co-activation during sensory stimulation. By comparing spontaneous and sensory-evoked events categorized as clustered or single, we find no intrinsic differences in amplitudes, emphasizing the role of co-activation in shaping event magnitude. Next, we optogenetically activated cerebellar nucleo-olivary (N-O) axons, a pathway central to theories of CS generation. To our surprise, while this robustly suppressed spontaneous IO activity, sensory-evoked events showed no reduction in either their probability or waveform. Our findings suggest that the traditional view of the N-O pathway as purely inhibitory or desynchronizing might be complemented with selective suppression of background activity while preserving sensory-driven responses. Together with the role of local co-activation in shaping IO event magnitudes, this work offers new insight into the timing and variation in complex spikes and their functional significance for behavior.

## Introduction

The field of cerebellar research is experiencing a renaissance. Not only is the role of cerebellar systems in non-motor behaviors increasingly well-accepted (***Van Overwalle et al. (2020); Hull (2020); Habas (2021); Kim et al. (2024); van der Heijden (2024)***), but a recent flurry of anatomical discoveries has unveiled complexities previously unappreciated. These include findings such as Purkinje neurons being innervated by multiple climbing fibers (***Busch and Hansel (2023)***) and directly targeting brainstem nuclei (***Chen et al. (2023)***), GABAergic cerebellar afferents and glutamatergic nucleoolivary axons (***Judd et al. (2021); Wang et al. (2023)***), and inhibitory projections to the inferior olive (IO) from additional brain regions beyond the cerebellar and vestibular nuclei (***van Hoogstraten et al. (2024)***). These observations challenge established views of olivo-cerebellar circuitry (e.g., ***Miall (2022); Carey (2024)***), suggesting a more intricate cerebellar network than previously recognized. Updating existing models of cerebellar circuitry to incorporate these findings is essential for gaining a comprehensive perspective on information processing within the olivo-cerebellar system and its computational interactions with other brain regions (***Zang and De Schutter (2023); Jun et al. (2024); de Xivry and Diedrichsen (2024)***).

However, amidst these advancements, most fundamental aspects of cerebellar function remain underexplored. Among these, the mechanisms determining the timing and properties of complex spikes in living animals remain largely unclear, despite being recognized as a cornerstone of cerebellar function since the early years of research. Moreover, despite recent appreciation of the distinct functional characteristics among cerebellar microzones (***De Zeeuw et al. (2021); De Zeeuw (2021); Blot et al. (2023)***), no systematic anatomical or functional distinctions have been identified between the olivary subnuclei—the principal and dorsal accessory olives—that target the hemispheric cerebellum.

This gap in knowledge extends to our understanding of the cerebellar feedback pathways to the IO. Traditionally, these pathways were thought to either suppress olivary activity—thereby contributing to cerebellar learning by diminishing “error” signaling—or desynchronize activity by uncoupling gap junctions between olivary neurons (***Llinas (1974); Rasmussen and Hesslow (2014); Lang et al. (2017); Lisberger (2021)***). Indirect evidence supporting the desynchronization of olivary activity by nucleo-olivary feedback has been reported (***Lang et al. (1996); Wagner et al. (2021)***). However, while experimental manipulation of IO activity or gap junctional coupling leads to expected effects in complex spike rate and synchrony (e.g., ***Lang et al. (1996); Blenkinsop and Lang (2006)***), evidence supporting the central notion that the nucleo-olivary pathway implements context-dependent suppression of sensory-related complex spikes (***Andersson et al. (1988); Ruigrok and Voogd (1995)*** but see ***Kim et al. (2020)*** has been limited. Importantly, our *in vitro* work (***Lefler et al. (2014)***) demonstrated only weak hyperpolarization of IO neurons upon optogenetic activation of the nucleoolivary axons. While even small hyperpolarizations can reduce the probability of isolated synaptic events generating olivary spikes (***Loyola et al. (2023)***), the excitability of IO neurons is dramatically lower in *in vitro* than *in vivo* and thus it remains uncertain to which extent the hyperpolarization caused by nucleo-olivary activation can effectively suppress incoming sensory signals in the intact brain.

One reason for the sparsity of reports is the technical difficulty of studying the IO in living animals, as it resides in the most ventral part of the caudal medulla, difficult to reach through the overlying brain structures. As a result, much of our knowledge of IO function comes from *in vitro* studies or indirect measurements of complex spike activity in the cerebellar cortex. While such studies have been instrumental in constructing current models of cerebellar function, they lack the context of intact neural circuitry and physiological conditions present *in vivo*. In particular, electrical coupling between neighboring IO neurons—mediated by gap junctions that are highly sensitive to physiological conditions difficult to replicate *in vitro*—has been proposed as a central element of olivary function (***Blenkinsop and Lang (2006); Placantonakis et al. (2006); Jacobson et al. (2008); De Zeeuw et al. (1998)***). Therefore, to fully comprehend the IO’s role and the nuances of its function, it is essential to observe its activity directly within the living brain, where olivo-cerebellar dynamics interact with gap junction-coupled olivary networks.

Here, we take advantage of a novel experimental preparation that allows calcium imaging-based monitoring of spontaneous and sensory-evoked activity in an anesthetized but intact mouse IO, together with localized activation of cerebellar nucleo-olivary axon terminals. This approach allows visualization and comparison of spontaneous activity in different IO subnuclei *in situ*, as well as direct examination of the modulation of IO activity by cerebellar feedback. Unexpectedly, we discovered that the modulation of IO events is significantly influenced by co-activation levels, revealing a previously unrecognized mechanism affecting complex spike properties.

## Results

### Mouse inferior olive is accessible for fluorescence imaging experiments through ventral side

In order to visualize network activity in a living mouse inferior olive (IO) and examine how it responds to activation of the nucleo-olivary (N-O) axons, we utilized a combination of custom-made and commercial viral vectors to transfect the IO and contralateral cerebellar nuclear (CN) neurons with GCamP6s and ChrimsonR, respectively (Figure 1A1-3; ***Dorgans et al. (2022)***). N-O neurons were transfected with ChrimsonR by AAV9-vector injection into the CN (Figure 1A1-2). For the IO, the injection parameters were tuned to optimize reliable transfection of neurons in the ventrolateral aspects of the principal and dorsal accessory olives (PO and DAO, respectively; Figure 1A3). After 2 - 3 weeks of transfection, the animals were prepared for the ventral aspect surgery (***Guo et al. (2021)***). Briefly, the skin, visceral tissue, and small parts of the ventral vertebra bone were removed in mice that were deeply anesthetized via tracheal cannula, providing clear visual access to the ventral surface of the medulla, allowing localization of the ventralmost aspects of PO and DAO based on vascularization (Figure 1B1-2). The exposed regions of the IO reside in the middle of the IO anterio-posterior extent, and correspond to the ventral bend of the dorsal leaf of the PO and the ventrolateral leaf of the DAO (indicated with shadowed regions in Figure 1A3). These subregions of PO and DAO send climbing fibers to the contralateral Crus 1 and Copula pyramis / Paramedian lobules, and collateralize mainly into the lateral and interposed cerebellar nuclei, respectively (e.g. ***Voogd et al. (2003); Sugihara and Quy (2007); Owusu-Mensah et al. (2023); Ruigrok and Voogd (2000); Sugihara (2011);*** Figure 1B3).

**Figure 1.**
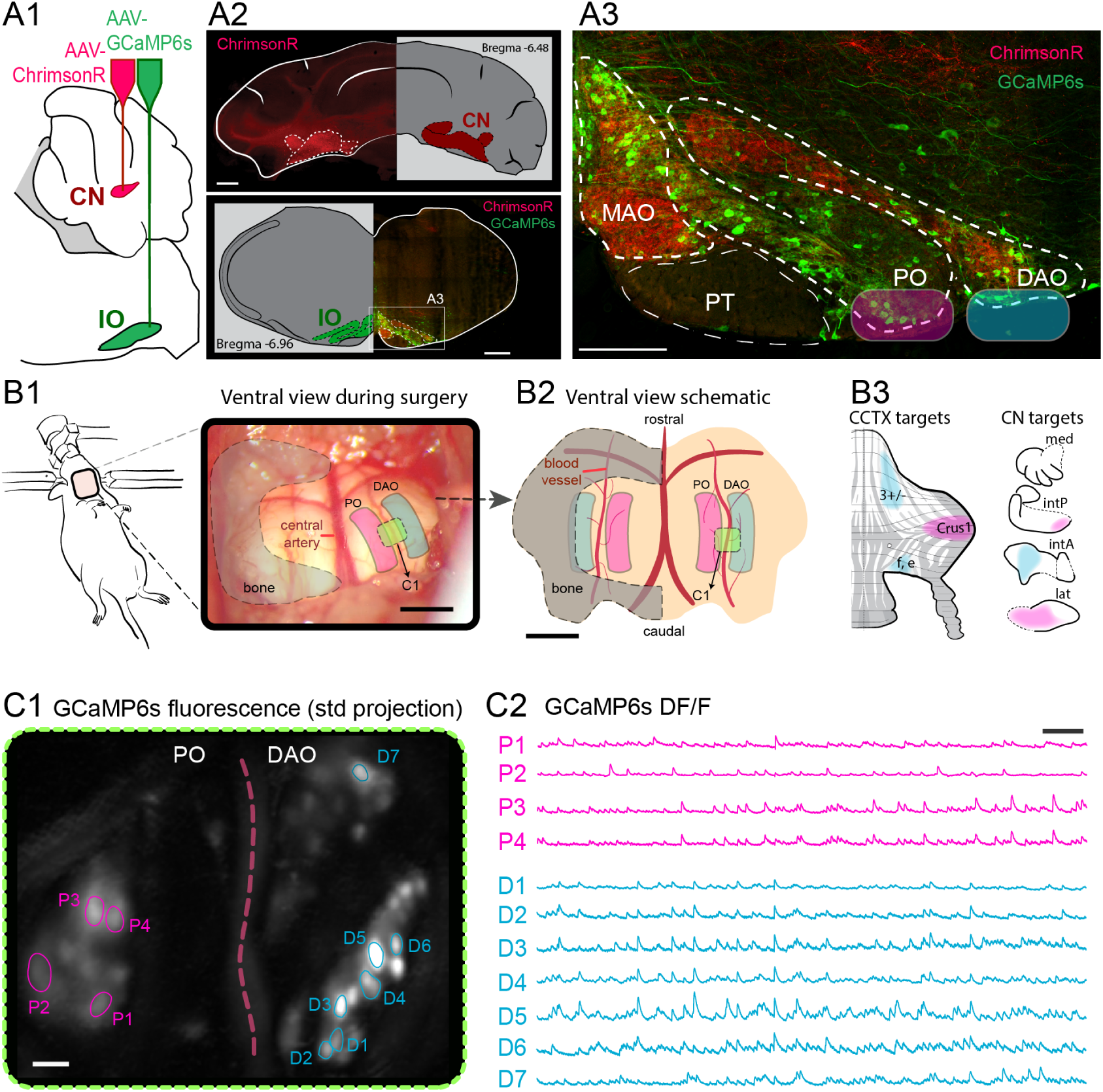
Imaging calcium events in living IO from the ventral side. (A1-3) Transfection of IO and CN neurons using AAV9-based viral vectors results in GCaMP6s expression in IO neurons and ChrimsonR in nucleo-olivary axons. (A1) Schematic of viral transfection targeting CN and IO. (A2) Confocal images of transfected CN (top) and IO (bottom) regions (10x maximum projection). (A3) Higher-resolution confocal image (20x) highlights transfected axons in the medial accessory olive (MAO), principal olive (PO), and dorsal accessory olive (DAO) with mediolateral imaging areas shaded (pink and cyan). (B1-3) Surgical access to the ventral medulla enables imaging of the IO. (B1) Ventral view during surgery. (B2) Schematic shows the ventral IO location, with key features like the blood vessel dividing PO and DAO marked. (B3) Schematic CCTX and CN regions targeted by PO and DAO climbing fibers. (C1-2) Example calcium recording from the IO. (C1) Standard deviation (STD) projection of a recording showing somata in PO (pink) and DAO (cyan), with manually marked regions of interest (ROIs). The red dashed line indicates the dividing blood vessel. (C2) Baseline-normalized (DF/F) intensity traces of fluorescence fluctuations in selected somata from PO (P1-P4) and DAO (D1-D7). Abbreviations: CN, cerebellar nuclei; IO, inferior olive; MAO, medial accessory olive; PT, pyramidal tract; PO, principal olive; DAO, dorsal accessory olive; CCTX, cerebellar cortex; med, intP, intA, lat: medial, posterior interpositus, anterior interpositus, lateral CN. Scale bars: 500 *μ* m (A2); 200 *μ* m (A3); 1 mm (B1-2); 20 *μ* m (C1); 20 s (C2). **Figure 1—video 1.** Spontaneous activity in PO and DAO.

Placing a 1 mm diameter 9 mm long GRIN lens (Inscopix, CA, USA) coupled to a miniature wholefield microscope on top of the exposed but intact *dura mater* of the medulla containing the inferior olive allowed capturing fluctuations of GCamP6s fluorescence (Figure 1C1-2). Although the dendrites are not well resolved in this recording fashion, the somata are clearly discernible and the presence of a blood vessel between the PO and the DAO allows unequivocal identification of the recording location. Crucially, careful placement of the GRIN lens across the dividing blood vessel allows simultaneous monitoring of events that lead to complex spikes in distant regions of the cerebellar cortex.

### Comparison of spontaneous activity in PO and DAO reveals modulation by co-activation

To explore the differences in activity in these two regions of IO that project to mainly zebrin positive (PO) and negative (DAO) parts of the cerebellar cortex, we combined GCamP6s recordings from 15 animals, totaling 20 and 16 recording sites in PO and DAO consisting of 121 and 72 neurons, respectively. Figure 2A shows example traces from a simultaneously recorded PO (pink) and DAO (cyan) neurons both as “raw DF/F” and “background-subtracted” forms (top and bottom in respective panels; see Methods for details of signal processing). In total, 2084 and 1176 spontaneous events were detected from DAO and PO neurons, with widely varying shapes, as expected for the broad range of IO spike waveforms (***Llinás and Yarom (1981); Mathy et al. (2009); Bazzigaluppi et al. (2012a); De Gruijl et al. (2012)***). To account for the experimental sample structure, non-normality, and potential confounders (e.g., random variation in GCaMP6s expression levels across neurons), we employed generalized linear mixed modeling (GLMM) for statistical comparisons throughout this study. Here, the analysis revealed that while the average time courses of calcium events were similar between PO and DAO neurons (Figure 2B1), events in PO reached larger values than those in DAO even though the average values did not differ (PO amplitude: 13.53 ± 8.61%, DAO amplitude: 11.47 ± 5.73%, *p*_GLMM_ = 0.20; PO FWHM: 1.50 ± 0.65*s*, DAO FWHM: 1.39 ± 0.66*s*, *p*_GLMM_ = 0.10; PO rise duration: 0.71 ± 0.24*s*, DAO rise duration: 0.69 ± 0.22*s*, *p*_GLMM_ = 0.22. All descriptive values here and in the following sections are presented as *mean* ± *SD*. Figure 2B2-3; see Tables 1, 2).

**Figure 2.**
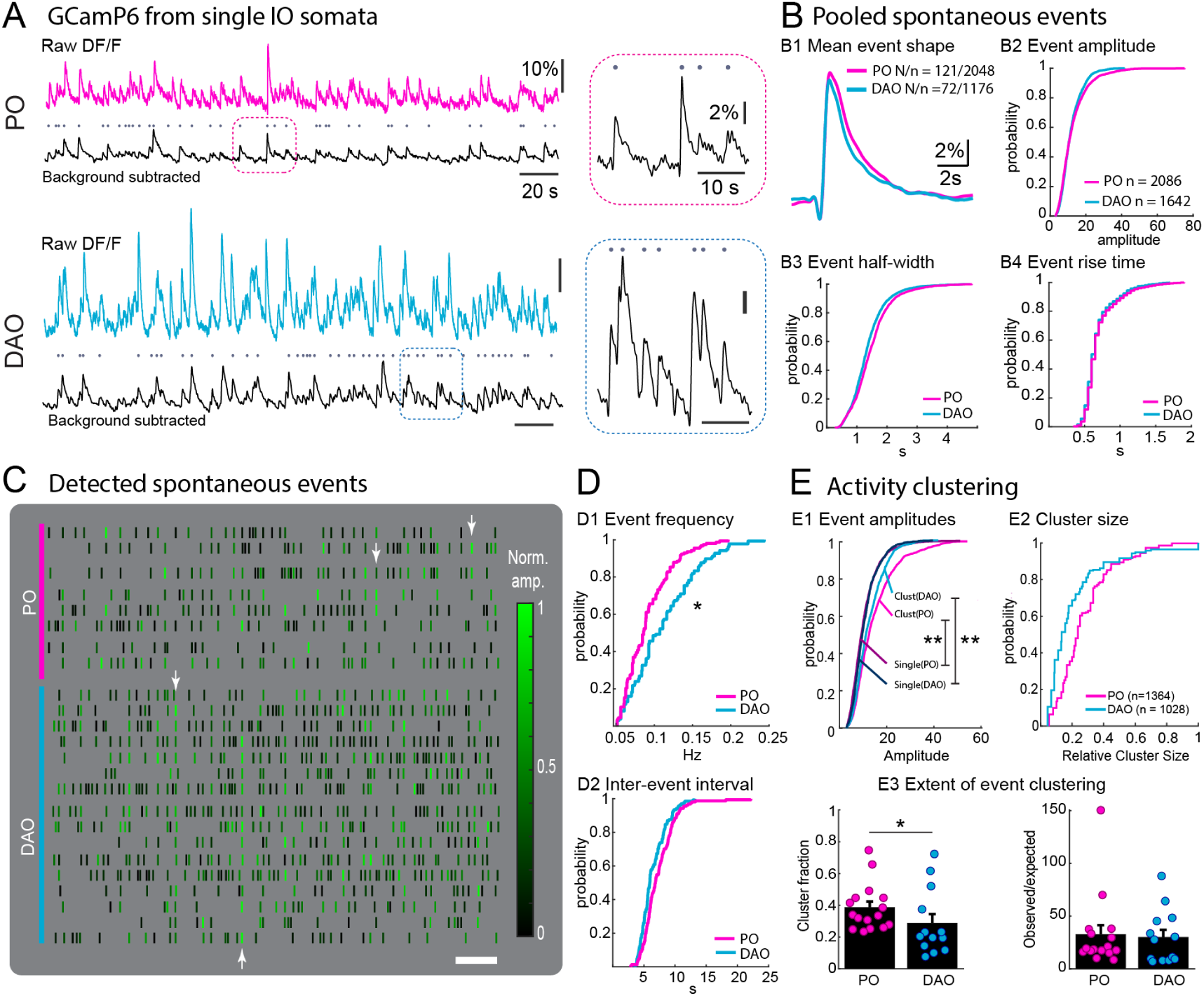
Spontaneous event properties in IO subnuclei display synchronicity-related modulation. A, example fluorescence imaging traces from a PO (pink) and DAO (cyan) neurons. Top traces display unprocessed DF/F values; bottom traces display the same data after background subtraction. Black dots denote events detected from the background-subtracted trace. See Methods for description of the processing. Regions of the background-subtracted traces indicated with dashed rectangles are shown with expanded scale insets to the right. B, summary of basic event shape characteristics. B1, average event shapes from PO and DAO neurons; shaded region denotes std. Events are vertically aligned at rise onset. B2, cumulative histogram of event amplitude distributions. Amplitudes are shown normalized to the baseline “noise” (see Methods for details). B3, empirical cumulative distribution of event widths (measured at half-amplitude). B4, empirical cumulative distribution plot of event rise times. C, spike raster visualization of the events detected in the recording shown in Figure 1. Each spike is color-coded by its amplitude, normalized to the largest event in a given cell. White arrowheads point to some instances when several large-amplitude events occur synchronously. Scale bar, 25 s. D, cumulative histogram of mean event frequency (D1) and the inter-event interval (D2). E, clustered event amplitude distributions show shift to larger values compared to asynchronous events both in PO and DAO recordings. E1, empirical cumulative distribution of clustered (bright colors) and single (dark colors) events in PO and DAO. Clustered spikes are defined with at least 2 neurons spiking within one frame (50 ms) of each other. For results with other definitions, see Figure 2 - Supplement 1. E2, cluster sizes (shown with empirical cumulative distribution plots relative to number of cells visible in a recording) are larger in PO than in DAO. E3, comparison of clustering between PO and DAO. Left, fraction of events recorded that were classified as clustered; right, normalized by expected value. Each dot represents a single field-of-view recording. Abbreviations: DF/F, fluorescence normalized by baseline intensity; PO, principal olive; DAO, dorsal accessory olive; N refers to number of cells, n to number of events. *, p < 0.05; **, p < 0.01. **Figure 2—figure supplement 1.** Event amplitude-width relationships **Figure 2—figure supplement 2.** Clustering effect on amplitude with various definitions of clustering.

**Table 1.**
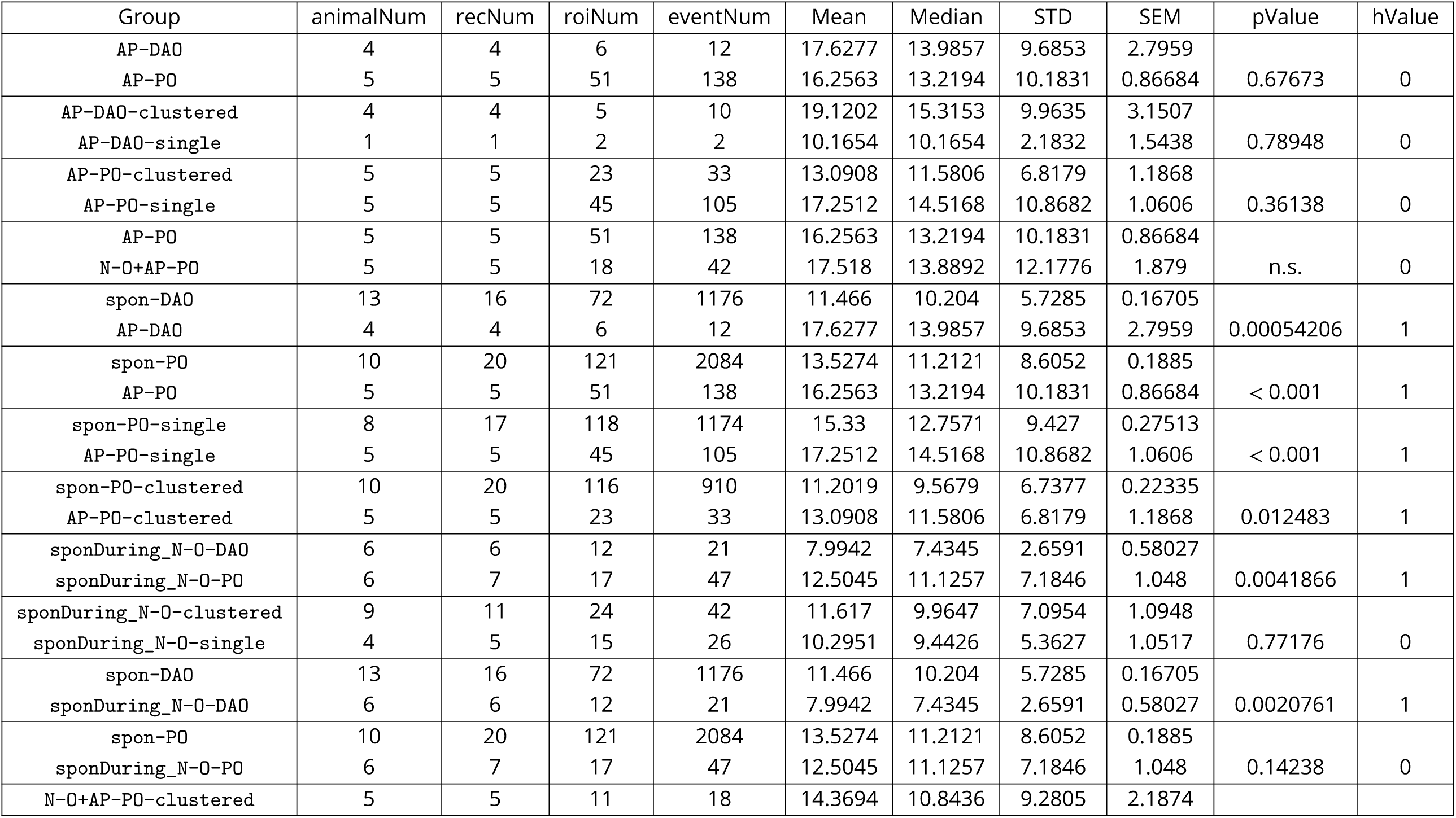
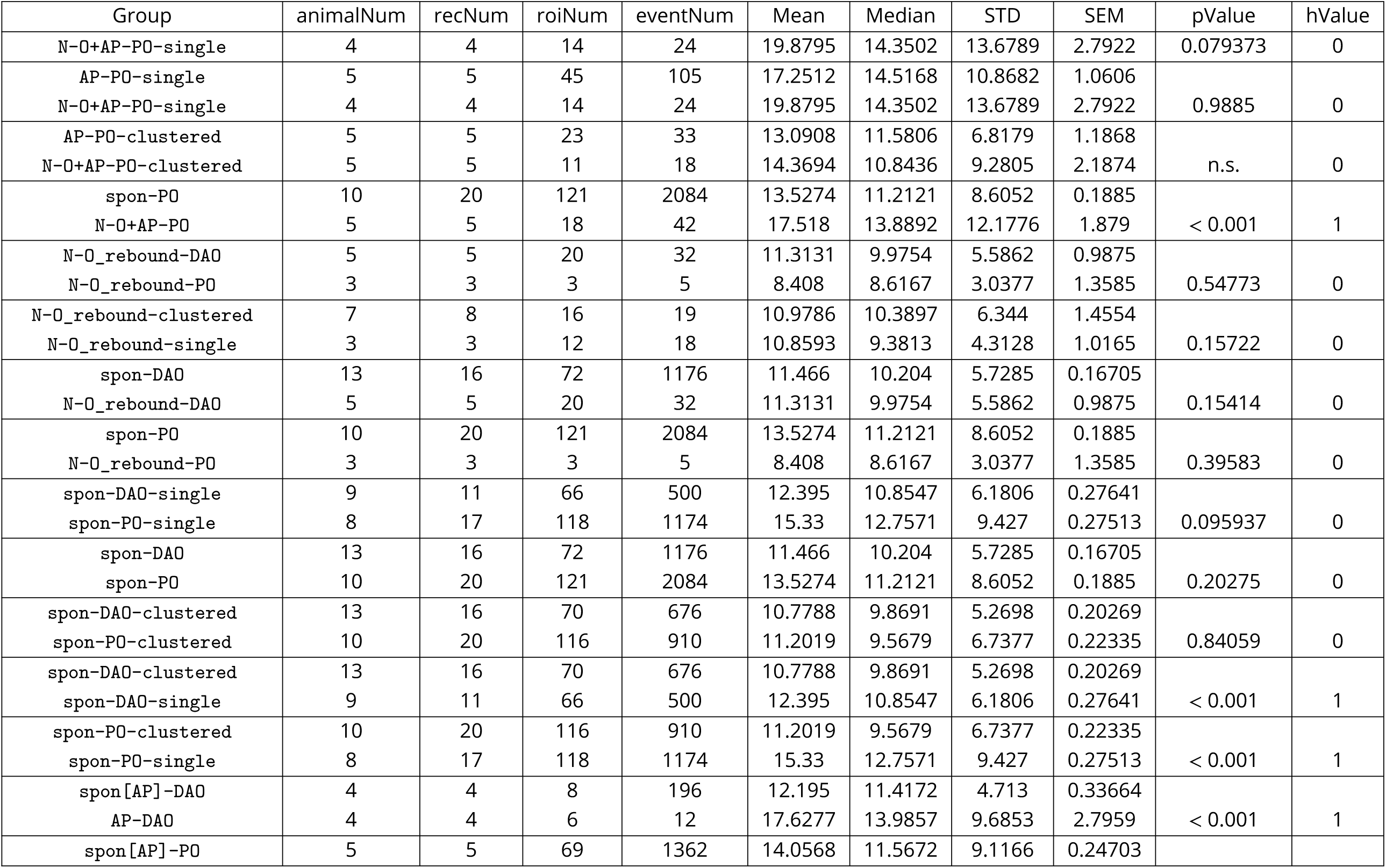
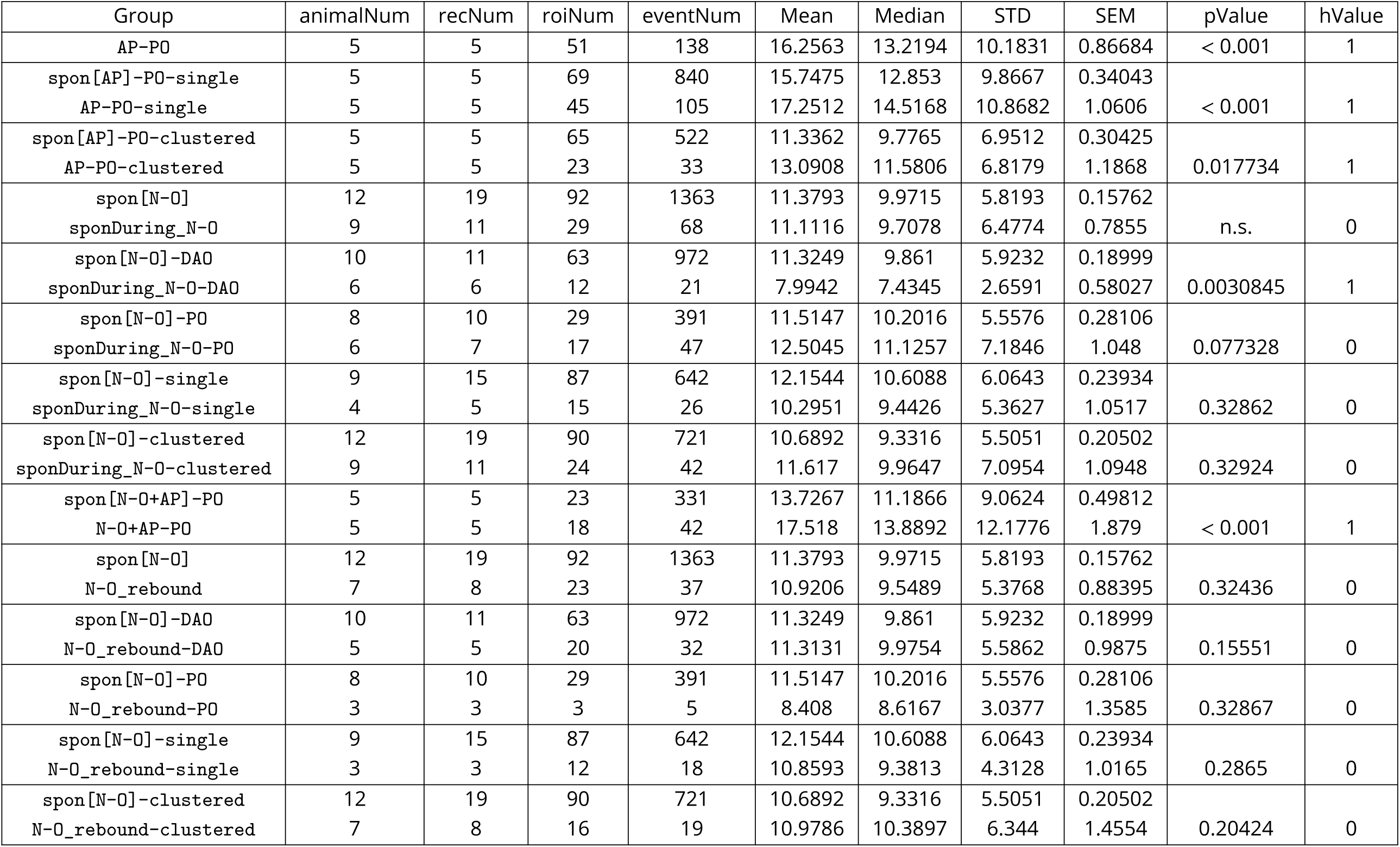
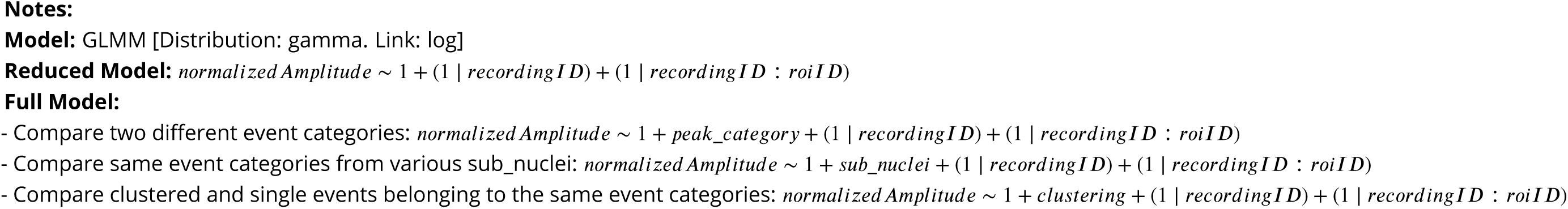
Normalized amplitude of calcium events. GLMM analysis.

**Table 2.**
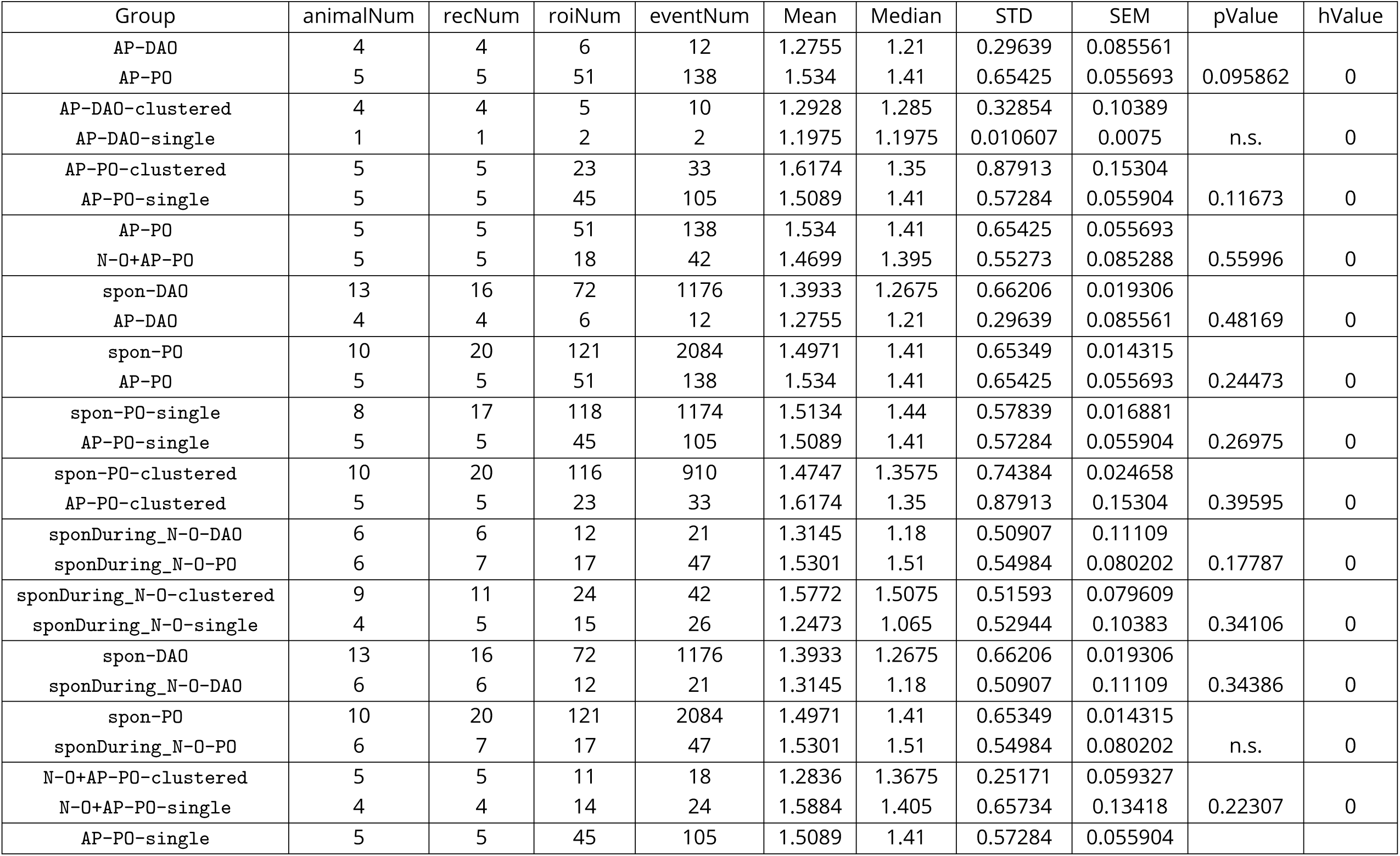
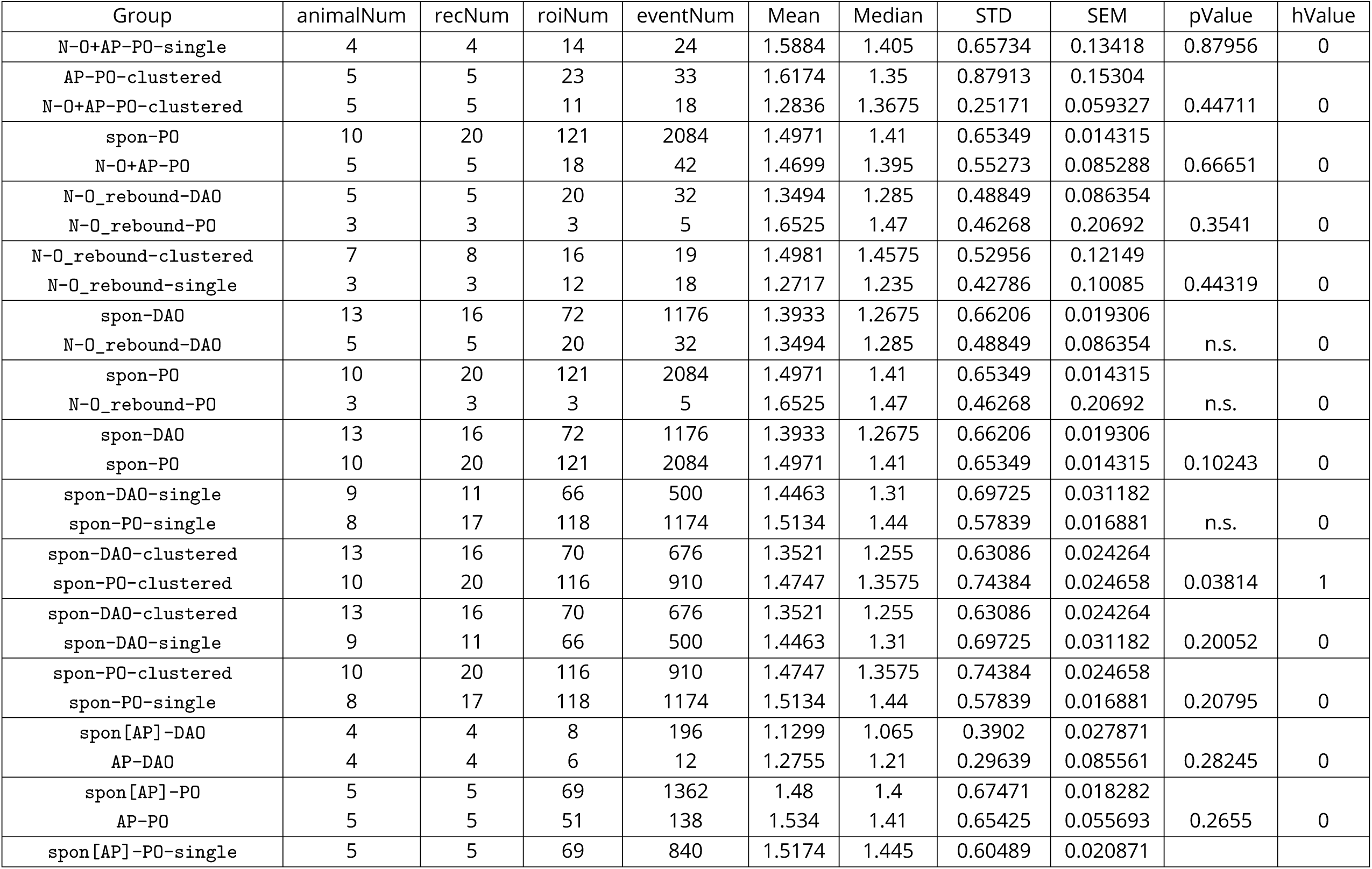
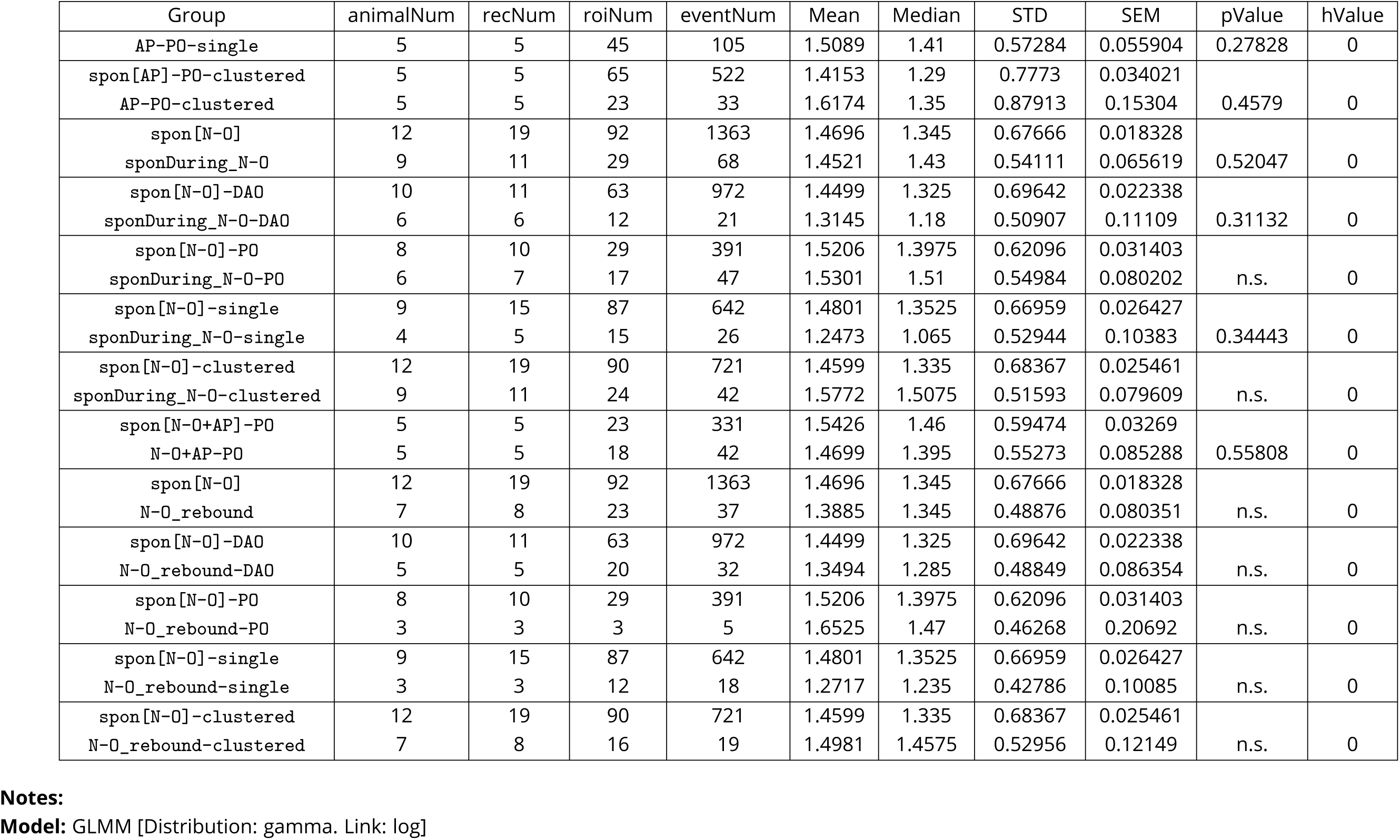
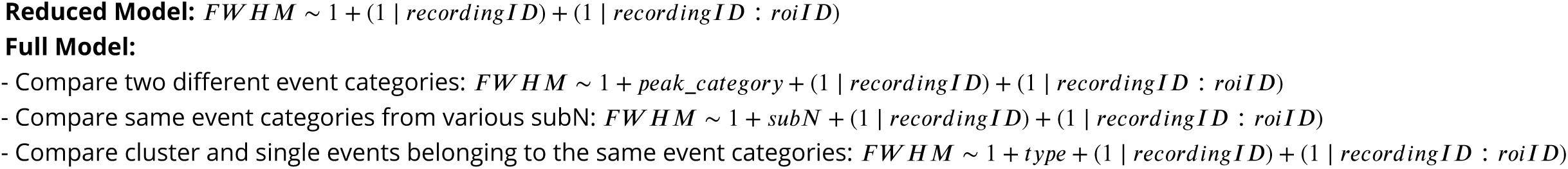
FWHM of calcium events (s). GLMM analysis.

The amplitudes and widths of the events showed a quasi-linear correlation (Figure 2 supplement 11) that flattened with the largest events, in line with our previous work showing that the dynamic range of the calcium indicator begins to saturate with the broadest IO spikes (> 10 ms; c.f. Figure 8C in ***Dorgans et al. (2022)***). Thus, even though with caution, we conclude that properties of the GCamp6s-generated events in our recording conditions can be informative of the duration of the underlying electrophysiological spike waveform.

Figure 2C shows activity in PO and DAO in spike raster form (same data as shown in Figure 1) with each event color-coded according to its amplitude. Due to the deep anesthesia, the average event frequency and inter-event interval were on average lower than previously reported for anesthetized mice and varied considerably within a single recording site (e.g. ***Bosman et al. (2010);*** PO frequency: 0.09 ± 0.03*Hz*, DAO frequency: 0.12 ± 0.04*Hz*, *p*_GLMM_ = 0.02; PO ISI: 1.50 ± 0.65*s*, DAO ISI: 6.68 ± 2.32*s*, *p*_GLMM_ = 0.35; PO rise duration: 7.69 ± 2.56*s*, DAO rise duration: 0.69 ± 0.22*s*, *p*_GLMM_ = 0.22. Figure 2D; see Table 3 for details of statistical descriptors.) The color-coded raster plot visualization revealed a surprising phenomenon: the spikes with largest amplitudes within a given cell occurred most often roughly simultaneously among several other cells (white arrowheads in 2C). This observation that “clustered” events (defined as being part of a group of at least two individual events in two cells with their peaks not more than one frame (50 ms) apart) are larger was consistently seen over the entire dataset: overall, clustered event amplitudes were on average around 40 and 18 % larger than in the “single” events (PO and DAO, respectively; PO event amplitudes (baseline normalized): clustered (15.33 ± 9.43%) vs single (11.20 ± 6.74%), *p*_GLMM_ < 0.001; DAO event amplitudes (baseline-normalized): clustered (12.40 ± 6.18%) vs single (10.78 +−5.27%), *p*_GLMM_ < 0.001. Figure 2E1); see Table 1 for all measurements and statistical descriptors, and Supplementary Table for GLMM modeling details)

**Table 3.**
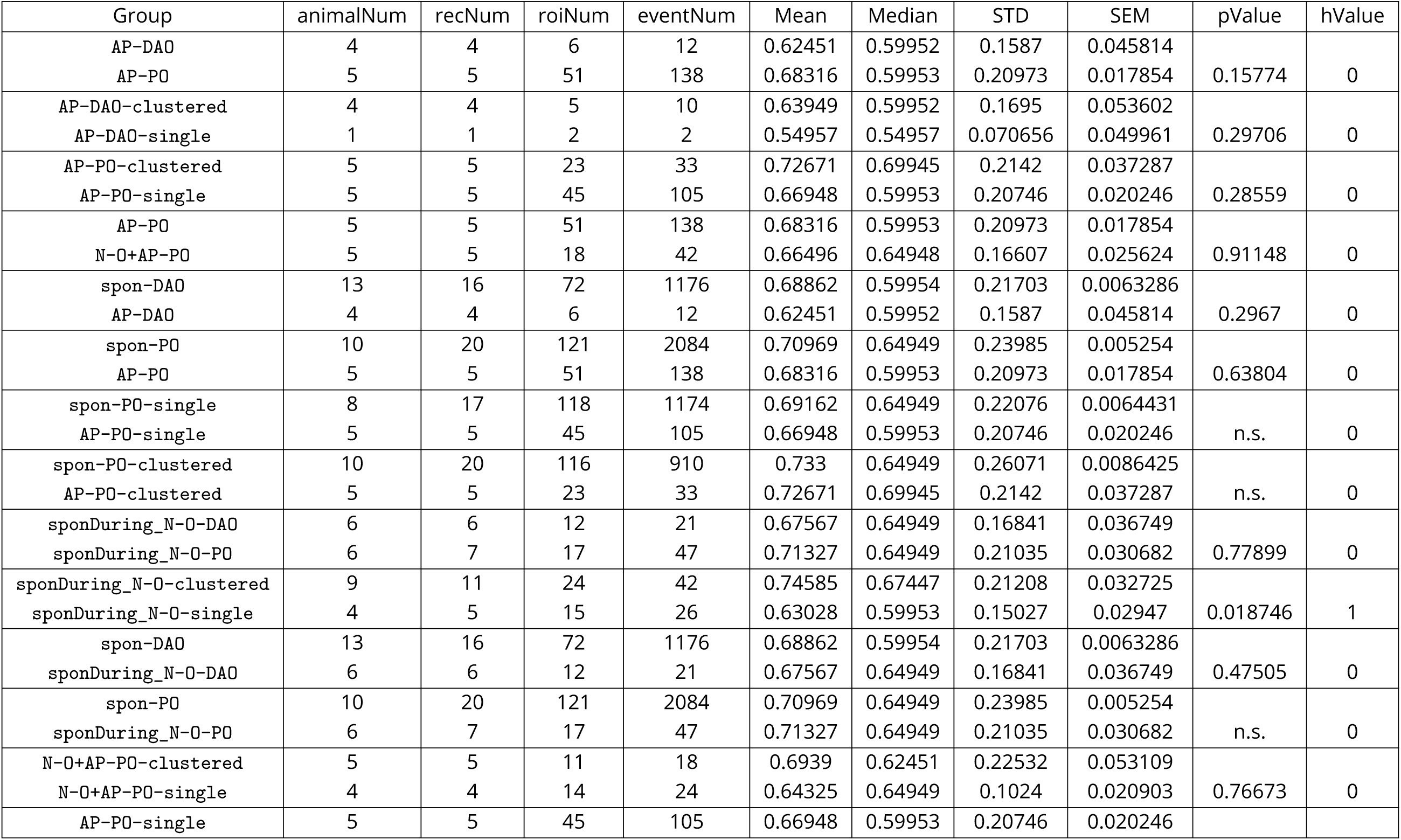
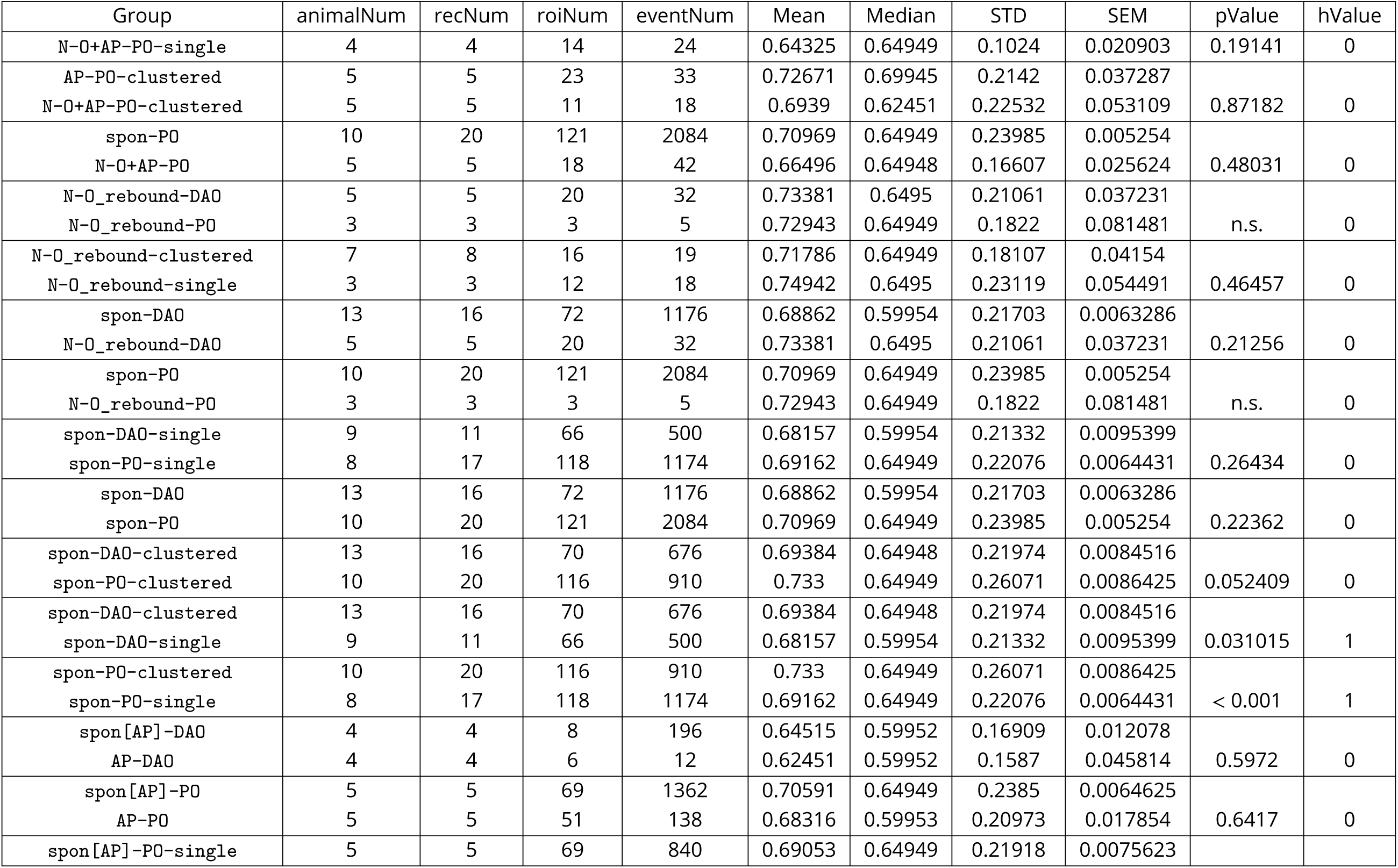
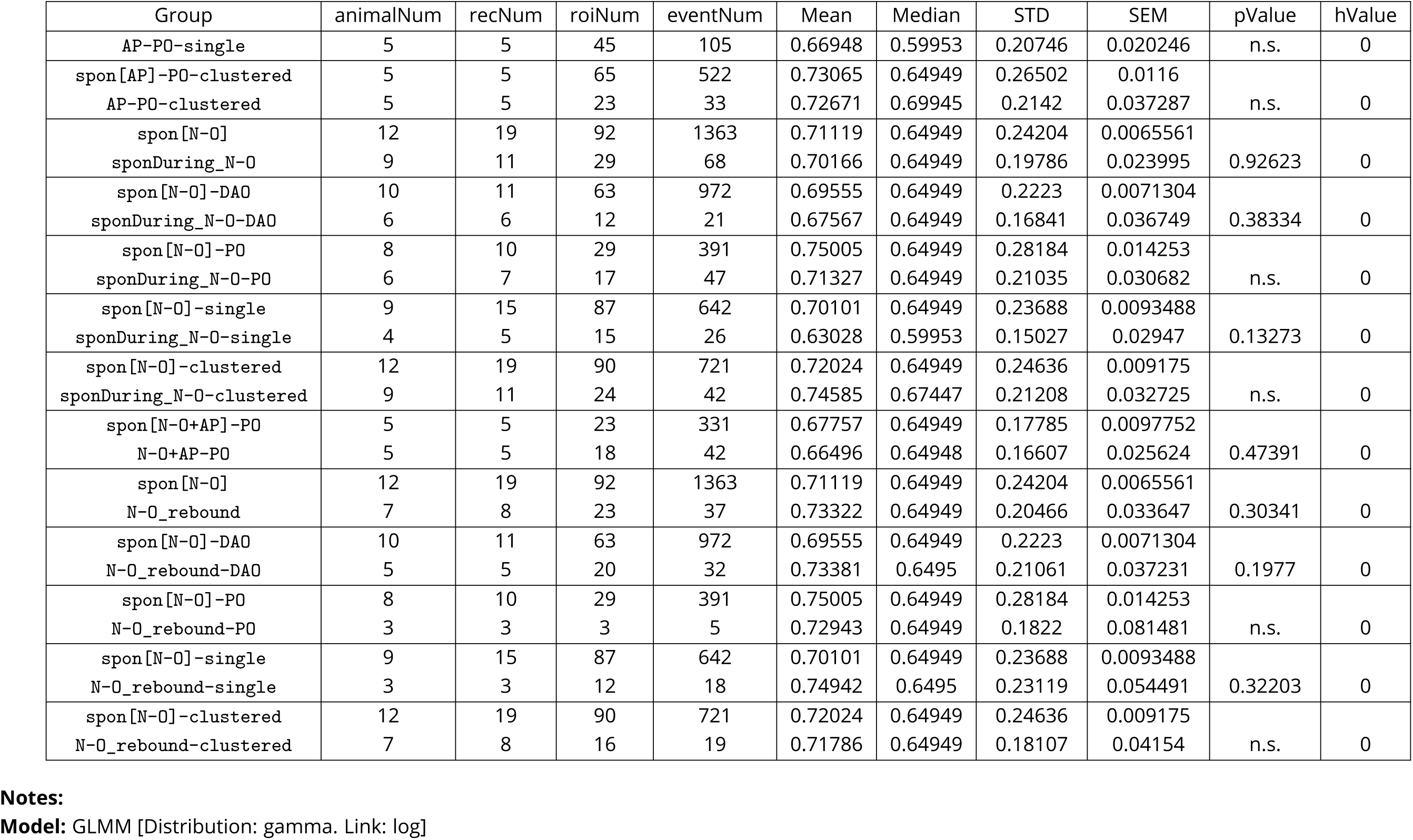
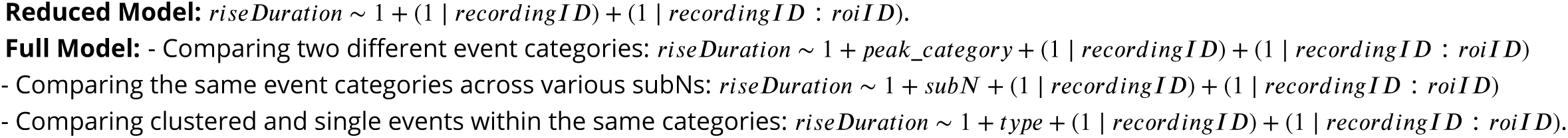
RiseDuration of calcium events (s). GLMM analysis.

Notably, the difference in amplitudes between “clustered” and “single” only became more pronounced when the criteria for clustering were made more strict by increasing the minimal number of active cells in an activity cluster and decreased when relaxing the temporal limits (Figure 2 supplement 2). Furthermore, when comparing the clustered and single events between PO and DAO it became apparent that the synchronicity seems to boost spike amplitudes more in PO than in DAO regardless of the definition of synchronicity 2F). This could be explained by the larger relative sizes of PO event clusters (Figure 2E2) leading to larger proportion of events being clustered in PO than DAO (Figure 2E3, left). However, when considering the higher event frequency of DAO, the difference in proportion of clustered events disappeared (Figure 2E3, right).

### Airpuff-evoked activity in principal olive

After establishing that spontaneous calcium event amplitudes in IO seem to increase with the amount of co-activation of nearby cells, we continued to examine if similar trends could be observed with events evoked by sensory stimulation. For this purpose, we chose to use a peri-ocular airpuff stimulation, as it could be argued to be the most commonly used one in the context of investigating the olivo-cerebellar system (***Yeo and Hesslow (1998); Heiney et al. (2021); Parras et al. (2022);*** Figure 3A1). While such stimulation has been reported to readily evoke complex spikes in cerebellar cortical regions targeted by both principal and dorsal accessory olives (***Sun (2012); Boele et al. (2010)***), among the recording sites used in this study, airpuff could reliably trigger events only in contralateral PO regions. As shown in Figure 3A2, the airpuff-evoked (AP) events were readily identifiable by being time-locked to the stimulation, and overall the AP delivery led to a roughly two-fold increase of event frequency (from 0.0960 ± 0.0431 Hz in the baseline period preceding stimulations to 0.2268 ± 0.2487 Hz, *p*_GLMM_ = 0.004; data from 5 mice, 69 cells and 631 stimulations; Figure 3A3; see TABLE 7 for details).

**Figure 3.**
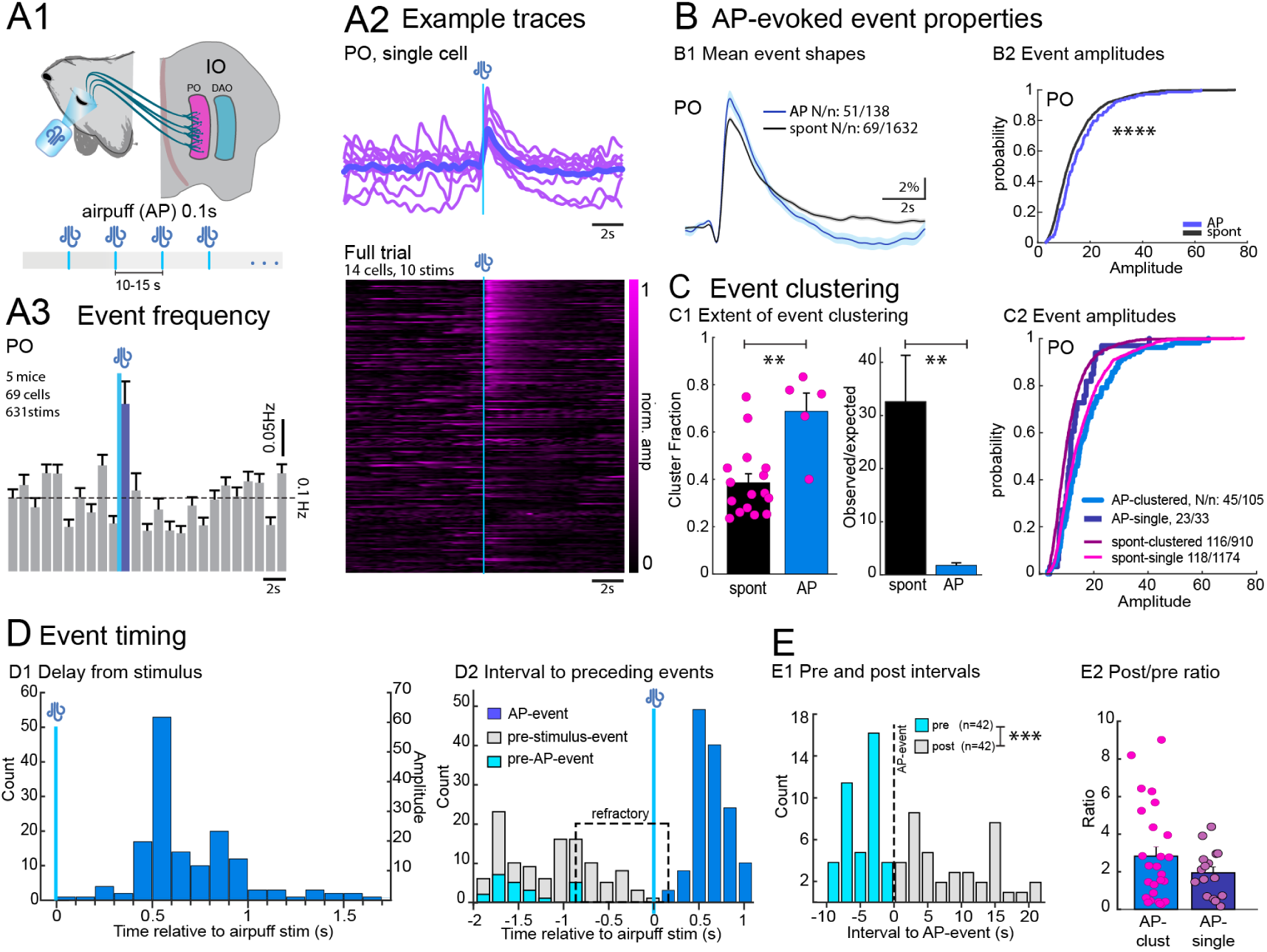
Airpuff-evoked events in the principal olive. Periocular airpuff stimulation (A1) reliably evoked events in PO neurons. (A2) Top: example traces from a single cell over 10 stimulations (dark trace: average). Bottom: heat-map of the full trial, with traces sorted by event amplitude. (A3) Peri-stimulus time histogram (mean ± SEM, 1-s bins) for all trials. B: AP-evoked event waveforms (B1) and amplitudes (B2) compared to the spontaneous events in the same cells and recordings. Waveforms are shown as mean ± SEM, vertically aligned at event onset. C, event clustering as defined for events shown in Figure 2. C1,(C) Event clustering. (C1) Clustering increased for AP-evoked events relative to spontaneous ones (left) but normalized to event rates (right). Note that for visual clarity markers are not shown on the right; see source data. C2, clustered and single AP-event amplitudes compared with spontaneous events from the same cells and trials. The difference between clustered and single AP-events does not reach significance with GLMM analysis due to low number of events per cell. D, AP event timing. D1, timing histogram of AP-evoked events with respect to stimulus. D2, timing histogram of AP-evoked events (blue) and preceding spontaneous events with respect to stimulus. Cyan: spontaneous events within trials with AP-evoked events, showing a refractory period > 1 s (dashed rectangle). (E) Intervals before and after AP-events. E1, comparison of intervals preceding (cyan) and following (gray) 42 AP-evoked events with a preceding and following spontaneous event within 10 s. E2, post/pre-interval ratios for clustered and single AP-events, with each marker representing a single stimulation. Abbreviations: IO, inferior olive; AP, airpuff; PO, principal olive; DAO, dorsal accessory olive; spont, spontaneous events; GLMM, generalized linear mixed model. **, p < 0.01; ***, p < 0.001; ****, p < 0.0001 **Figure 3—video 1.** AP-evoked activity in PO.

**Table 4.** Frequency of spontaneous calcium events (Hz). GLMM analysis.

| Group | animalNum | recNum | roiNum | Mean | Median | STD | SEM | pValue | hValue |
| --- | --- | --- | --- | --- | --- | --- | --- | --- | --- |
| spon-DAO | 13 | 16 | 72 | 0.11745 | 0.11217 | 0.042887 | 0.0050543 | 0.020221 | 1 |
| spon-PO | 10 | 20 | 121 | 0.090146 | 0.081116 | 0.033465 | 0.0030423 |  |  |

**Notes:**

**Table 5.**
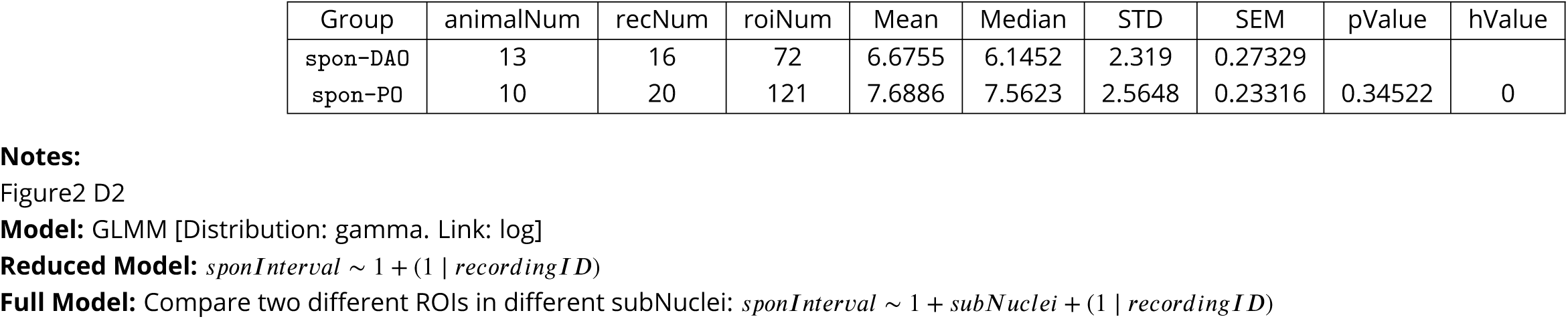
Interval of spontaneous calcium events (s). GLMM analysis.

| Group | animalNum | recNum | roiNum | Mean | Median | STD | SEM | pValue | hValue |
| --- | --- | --- | --- | --- | --- | --- | --- | --- | --- |
| spon-DAO | 13 | 16 | 72 | 6.6755 | 6.1452 | 2.319 | 0.27329 | 0.34522 | 0 |
| spon-PO | 10 | 20 | 121 | 7.6886 | 7.5623 | 2.5648 | 0.23316 |  |  |

**Notes:**

**Table 6.** Cluster fold PO vs. DAO. Mann-Whitney U test.

| Group | nNum | Mean | Median | STD | SEM | pValue | hValue |
| --- | --- | --- | --- | --- | --- | --- | --- |
| PO | 16 | 0.37541 | 0.34046 | 0.15244 | 0.038111 | 0.035748 | 1 |
| DAO | 14 | 0.2862 | 0.21165 | 0.19931 | 0.053267 |  |  |

**Notes:**

**Table 7.** Peristimulus (AP) event frequency in PO neurons (Hz). BootStrap analysis.

| Group | roiNum | Mean | Median | STD | SEM | pValue | hValue |
| --- | --- | --- | --- | --- | --- | --- | --- |
| baseline (6 bins pre-AP stimulation) | 69 | 0.0960 | 0.1000 | 0.0431 | 0.0052 | 0.004 | 1 |
| AP bin | 69 | 0.2268 | 0.1667 | 0.2487 | 0.0299 |  |  |

**Notes:** Figure3 A3 N-O bins were compared to the baseline

**Table 8.** Cluster fraction spon vs AP. Mann-Whitney U test.

| Group | nNum | Mean | Median | STD | SEM | pValue | hValue |
| --- | --- | --- | --- | --- | --- | --- | --- |
| spon | 16 | 0.37541 | 0.34046 | 0.15244 | 0.038111 | 0.0056719 | 1 |
| AP | 5 | 0.69113 | 0.76119 | 0.17537 | 0.078426 |  |  |

**Notes:** Figure3 C1 fraction

**Table 9.** Cluster fold spon vs AP. Mann-Whitney U test.

| Group | nNum | Mean | Median | STD | SEM | pValue | hValue |
| --- | --- | --- | --- | --- | --- | --- | --- |
| spon | 16 | 32.4231 | 19.0414 | 34.7958 | 8.699 | 0.0011078 | 1 |
| AP | 5 | 1.8285 | 2.2619 | 1.0106 | 0.45194 |  |  |

**Notes:** Figure3 C1 fold

**Table 10.**
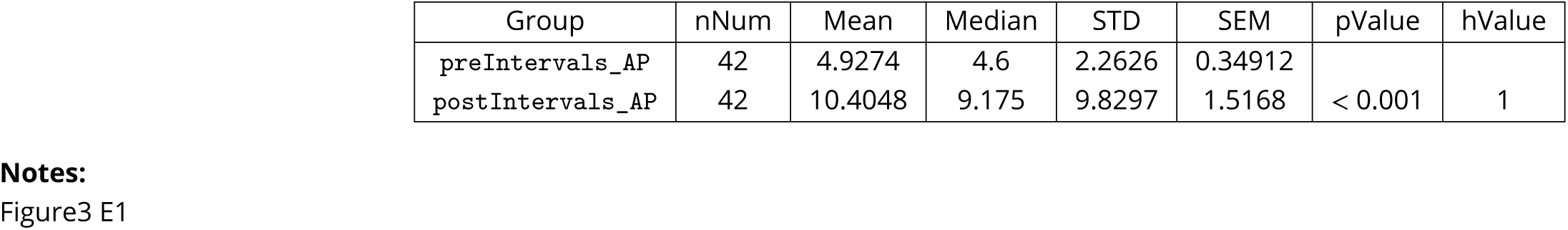
Pre and post intervals of AP events (s). Wilcoxon paired rank sum.

| Group | nNum | Mean | Median | STD | SEM | pValue | hValue |
| --- | --- | --- | --- | --- | --- | --- | --- |
| preIntervals_AP | 42 | 4.9274 | 4.6 | 2.2626 | 0.34912 | < 0.001 | 1 |
| postIntervals_AP | 42 | 10.4048 | 9.175 | 9.8297 | 1.5168 |  |  |

**Notes:**

**Table 11.** Pre and post ratios of clustered and single AP events. Mann-Whitney U test.

| Group | nNum | Mean | Median | STD | SEM | pValue | hValue |
| --- | --- | --- | --- | --- | --- | --- | --- |
| AP-clustered | 26 | 2.8269 | 2.0731 | 2.5583 | 0.50171 | 0.65036 | 0 |
| AP-single | 16 | 1.9381 | 1.8896 | 1.2583 | 0.31458 |  |  |

**Notes:**

**Table 12.** Peristimulus (N-O) event frequency in PO neurons (Hz). BootStrap analysis.

| Group | nNum | Mean | Median | STD | SEM | pValue | hValue |
| --- | --- | --- | --- | --- | --- | --- | --- |
| baseline (6 bins pre-N-O stimulation) | 28 | 0.0875 | 0.0833 | 0.0455 | 0.0086 |  |  |
| N-O bin-2 | 29 | 0.0560 | 0 | 0.1185 | 0.0220 | 0.012 | 1 |
| N-O bin-3 | 29 | 0.0359 | 0 | 0.0761 | 0.0141 | < 0.001 | 1 |
| N-O bin-4 | 29 | 0.0819 | 0 | 0.1262 | 0.0234 | 0.028 | 1 |
| N-O bin-5 | 29 | 0.1092 | 0 | 0.1370 | 0.0254 | 0.616 | 0 |

**Notes:**

**Table 13.**
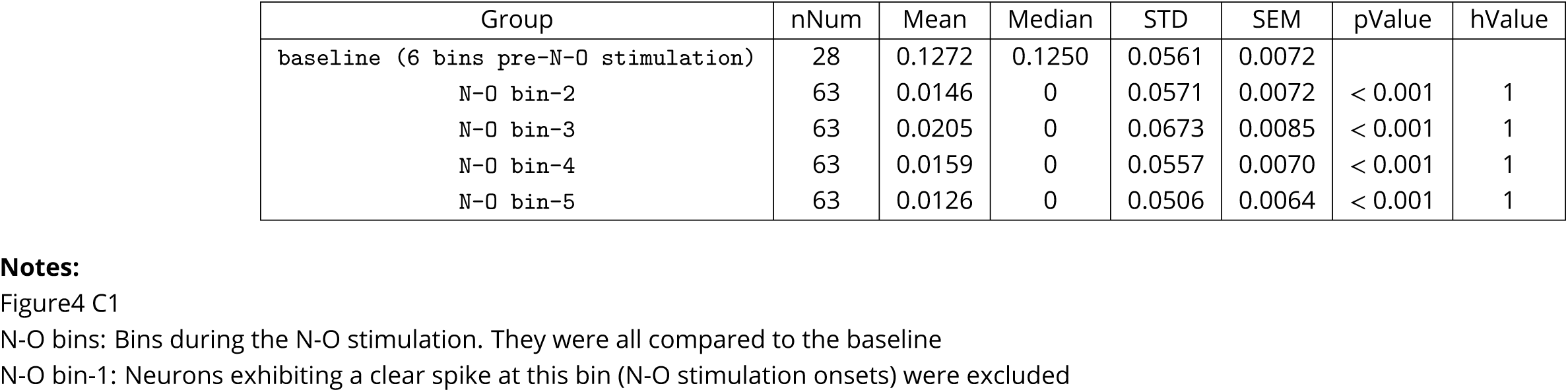
Peristimulus (N-O) event frequency in DAO neurons (Hz). BootStrap analysis.

| Group | nNum | Mean | Median | STD | SEM | pValue | hValue |
| --- | --- | --- | --- | --- | --- | --- | --- |
| baseline (6 bins pre-N-O stimulation) | 28 | 0.1272 | 0.1250 | 0.0561 | 0.0072 |  |  |
| N-O bin-2 | 63 | 0.0146 | 0 | 0.0571 | 0.0072 | < 0.001 | 1 |
| N-O bin-3 | 63 | 0.0205 | 0 | 0.0673 | 0.0085 | < 0.001 | 1 |
| N-O bin-4 | 63 | 0.0159 | 0 | 0.0557 | 0.0070 | < 0.001 | 1 |
| N-O bin-5 | 63 | 0.0126 | 0 | 0.0506 | 0.0064 | < 0.001 | 1 |

**Notes:**

In line with earlier works comparing sensory-evoked and spontaneous complex spikes recorded in the cerebellar cortex (e.g. ***Roh et al. (2020)***), the AP-evoked events were significantly larger than spontaneous events recorded in the same cells and trials (spontaneous: 13.53 ± 8.61%, AP-evoked: 16.26 ± 10.18%, *p*_GLMM_ < 0.001; Figure 3B1-2). Furthermore, the AP-evoked events occurred in temporal clusters more commonly than the spontaneous events (spontaneous: 0.38 ± 0.15, AP-evoked: 0.69 ± 0.18*Hz*, *p*_GLMM_ = 0.006; Figure 3C1, left). However, when the much higher event rate was accounted for, the extent of clustering was less impressive than for spontaneous activity (spontaneous: 32.42 ± 34.80, AP-evoked: 1.83 ± 1.01*Hz*, *p*_GLMM_ = 0.001; Figure 3C1, right), suggesting that the mechanism of temporal clustering could be different between spontaneous and evoked events. Nevertheless, the evoked event amplitudes showed similar modulation with clustering (Figure 3C2), even though the significance did not emerge from GLMM testing due to the much lower numbers of events (AP-clustered (17.25 ± 10.87%) vs AP-single (13.09 ± 6.82%), *p*_GLMM_ = 0.36. see TABLE 1).

### Timing of peri-stimulatory events in the principal olive neurons

One of the puzzling aspects of the IO function relates to its stubbornly invariant spiking rate. To our knowledge, no naturalistic manipulations of the experimental mice have been reported to elevate the average cerebellar CS rates beyond 1-2 Hz (***Negrello et al. (2019)***) even though stereotypical CS “doublets” do occur (***Titley et al. (2019)***).

Closer examination of the event timing with respect to AP delivery revealed a two-peaked distribution of the delays from the stimulation (Figure 3D1). However, the peaks did not correspond to occurrences of the same cell spiking twice. In our dataset, the shortest interval between a stimulation-evoked event and any preceding event was 1 s (Figure 3D1, cyan data) even though spontaneous events were seen in a considerable number of trials preceding the AP-stimulation by much shorter intervals (Figure 3D1, gray data), suggestive of a refractory period for successful AP-event generation.

Noting that in the post-stimulation period, stimulation-triggered average traces (such as shown in Figure 3A2) event frequency seemed to decrease and the event frequency was mildly lower (Figure 3A3, bars below dashed line), we wondered whether a post-stimulatory event rate suppression could be seen. As shown in Figure 3E1 for AP-evoked events with a spontaneous event detected within the preceding 10-s window (n = 42), the intervals to the following events were longer (preIntervals_AP: 4.93 ± 2.26*s*, postIntervals_AP: 10.40 ± 9.83*s*, *p*_Wilcoxon_ < 0.001). Curiously, the largest post/pre interval ratios were seen with events that were clustered (Figure 3E2), but on average the difference between clustered and single AP-events was not significant.

Despite the reduced excitability in anesthetized mice that complicates behavioral comparisons, periocular airpuff stimulation consistently activated PO neurons. This suggests that our ventral-approach methodology offers a powerful alternative to cerebellar cortical recordings when investigating the timing of complex spikes by mechanisms intrinsic to the IO.

### Optogenetic activation of nucleo-olivary axons suppresses spontaneous spiking in the IO

To examine the manner in which the cerebellum can affect complex spike generation via activity in the nucleo-olivary (NO) pathway, we used animals in which the red-shifted excitatory opsin ChrmisonR was expressed in the nucleo-olivary (N-O) axon terminals. 5-second trains of LED light (620 ± 30 nm, 20 Hz, 5 ms pulse) were delivered through the GRIN lens to the recording area for targeted activation following the insights obtained in earlier *in vitro* experiments (***Lefler et al. (2014);*** Figure 4A1)). Not surprisingly, optogenetic activation of N-O axons resulted in a decrease in spontaneous activity in both PO and DAO neurons (see Figure 4B1 for example calcium traces from single neurons, and B2 for compiled visualization of all stimulations in all neurons in single recordings). The suppression of spiking was most pronounced during the early part of the stimulation (Figure 4C1), with moderate recovery of spiking from 2 seconds onwards.

**Figure 4.**
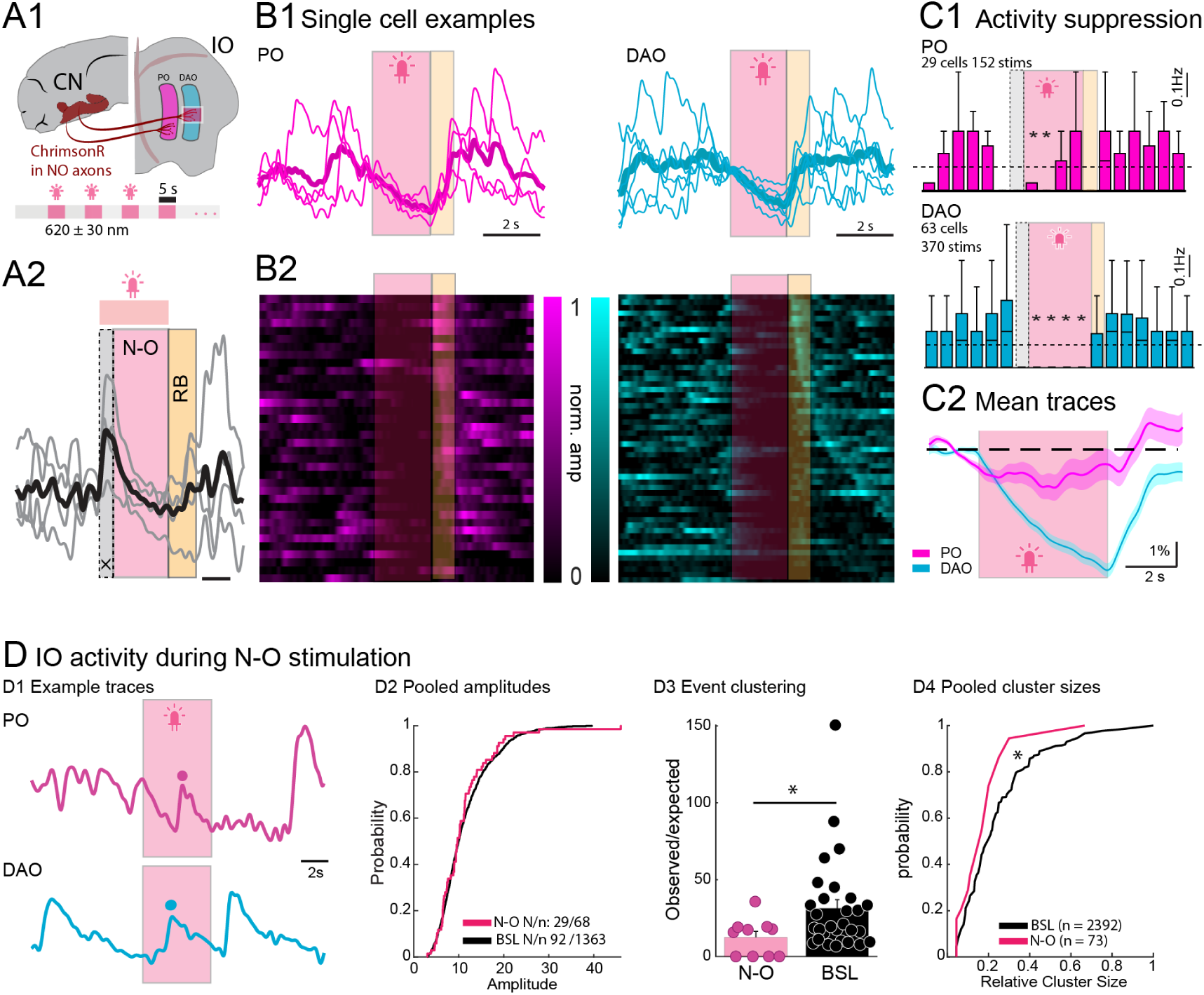
Changes in IO activity evoked by stimulation of N-O axons. (A) Experimental arrangement schematic (A1) and spike classification based on timing (A2). The first second of the stimulation period (gray box) is excluded from analysis; the yellow box denotes the rebound activity window. (B) Examples of NO stimulation effects on IO spiking in single cells (B1) and across all cells and stimulations in a single recording (B2), sorted by the time of the first spike after stimulus ends. In B2, color indicates event amplitude normalized to each cell’s maximum; shading corresponds to A2. Shading as in A2. (C) Summary of N-O stimulation effects on spike frequency in PO (C1, top) and DAO (C1, bottom) cells; box plots show median±SEM values. (C2) Mean±SEM of GCaMP6s fluorescence traces from all stimulation trials. (D) Example traces from PO (top) and DAO (bottom) cells with events during NO stimulation (shaded area) (D1). (D2) Cumulative distributions of event amplitudes during N-O stimulation (red) versus outside stimulation (black) in the same cells and trials; PO and DAO data are pooled due to low event numbers. (D3) Comparison of clustering extent, shown as the ratio of observed to expected clustered event fractions, for events during N-O stimulation (red) and outside stimulation (black). (D4) Cumulative distributions of cluster sizes during and outside N-O stimulation, relative to the total available cell pool. Abbreviations: CN, cerebellar nuclei; IO, inferior olive; N-O, nucleo-olivary; stims, stimulations; BSL, baseline (periods outside stimulation). Statistical significance indicted with asterisks as follows : *, p < 0.05. **Figure 4—video 1.** Effect of N-O stimulation in PO. **Figure 4—video 2.** Effect of N-O stimulation in DAO. **Figure 4—figure supplement 1.** IO events evoked by N-O stimulation. **Figure 4—figure supplement 2.** Activity after N-O stimulation.

Importantly, while most of these axons are GABAergic, recent work has highlighted the presence of glutamatergic CN neurons that project to the IO, especially to the medial accessory olive (MAO; ***Judd et al. (2021); Wang et al. (2023)***). As our recording sites were exclusively in the PO and DAO regions, we were surprised to find that in a significant fraction of recorded neurons (46 and 41 % of PO and DAO neurons) a spike was observed at the onset of N-O stimulation (Figure 4A2). These spikes were indistinguishable in shape between PO and DAO cells, and also between N-O-triggered and spontaneous events (see Figure 4, supplement 1). Due to our experimental conditions and the nonspecific viral targeting of ChrimsonR, we cannot be entirely certain that only GABAergic (inhibitory) N-O axons were transfected within the IO. To address this potential confounding effect, we chose to exclude neurons that exhibited a clear spike at the onset of N-O stimulation, as such spikes could indicate direct excitation rather than the intended inhibitory effect. Furthermore, to avoid potential contamination in our event frequency comparisons, we excluded the first second of the NO-stimulation period from our statistical analyses, even if no clear N-O-linked events were seen.

Overall, we detected 47 events in PO cells and 21 events in DAO cells during periods of NO-stimulation, excluding those that took place during the initial second (refer to Figure 4D1 for illustrative examples). The event amplitudes, compared to spontaneous events in the same neurons, were indistinguishable and no differences could be discerned between events seen in PO and DAO neurons (See TABLE 14; Figure 4D2). Consistent with the idea that N-O stimulation is expected to decrease the synchrony of IO activity, there was a noticeable decrease in both the percentage of clustered events when adjusted for event frequency (N-O: 11.48±12.15, BSL: 34.39±32.72, *p*_Mann-Whitney_ = 0.019; Figure 4D3) and the average cluster sizes (Figure 4D4). Similarly to spontaneous events, the amplitude of events observed during N-O stimulation was higher when clustered (Figure 4, supplement 2), but the effect did not reach significance, probably due to the very low number of events spread over numerous recordings (see TABLE 1).

**Table 14.**
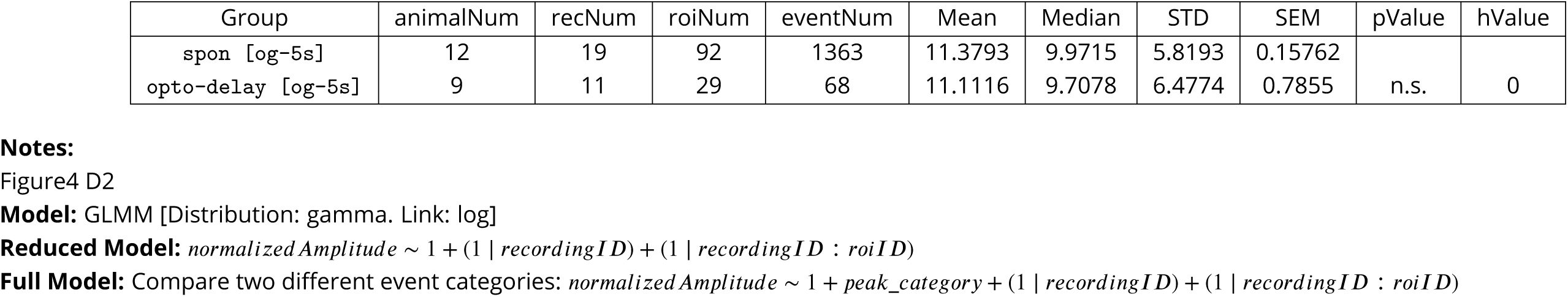
Normalized amplitude of events during N-O stimulation versus outside stimulation. GLMM analysis.

**Table 15.**
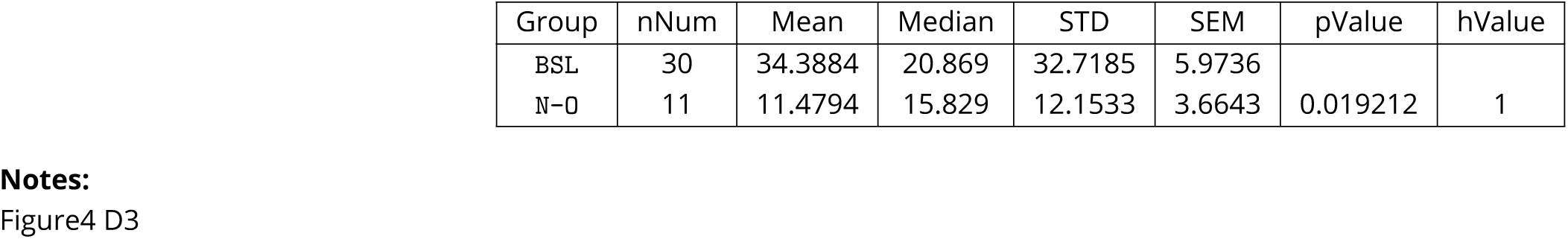
Cluster fold of spon and spon during N-O stimulation. Mann-Whitney U test.

**Table 16.**
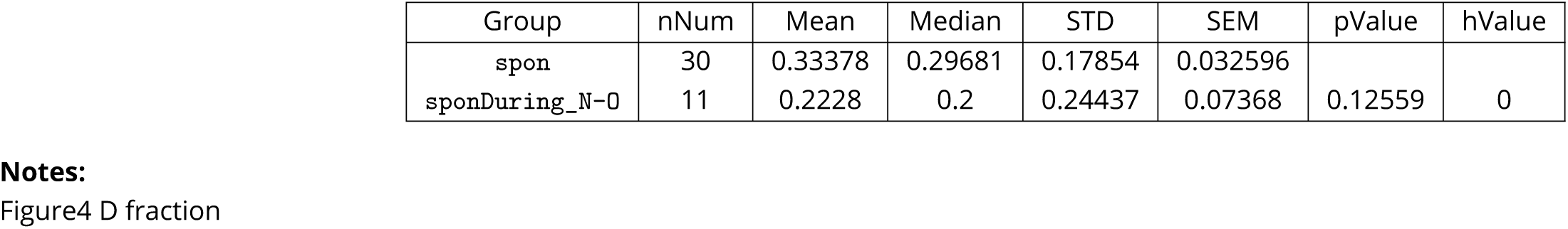
Cluster fraction of spon and spon during N-O stimulation. Mann-Whitney U test.

We also analyzed the behavior of IO neurons in the period just after the optogenetic light was deactivated (illustrated by the yellow shading in Figure 4 4A-C), to find evidence supporting the hypothesis that removing N-O-mediated inhibition triggers a “rebound spike” in IO neurons. Such rebound spikes have occasionally been reported when IO neurons are hyperpolarized *in vitro* by current injection but not with isolated stimulation of N-O axons (***Lefler et al. (2014); Loyola et al. (2023)***). In our experiments, a spike was sometimes detected within the 1-second window after the offset of N-O stimulation in a subset of neurons (5 events in 10% of PO neurons and 32 events in 32% of DAO neurons (Figure 4 supplement 2). However, these post-stimulation events did not differ in shape or clustering propensity from spontaneous events recorded in the same cells. Additionally, we found no correlation between the degree of inhibition during N-O stimulation and the probability of rebound activity in the post-stimulation window. Examining the timing of poststimulation events also did not reveal consistent delays between offset of stimulation and spike occurrence. Consequently, our findings do not validate the hypothesis that GABAergic N-O axons alone can induce a precisely-timed rebound spike in IO neurons. Additional investigations are necessary to assess if the excitability of IO neurons is increased following the N-O stimulation.

### Preferential suppression of spontaneous events by nucleo-olivary activity

Although activation of NO axons resulted in a clear decrease in spontaneous spike frequency in both PO and DAO, the definition of neuronal inhibition as reducing the probability of spike generation in response to a known input called for further experiments. Thus, we combined the optogenetic activation of NO axons with airpuff stimulations (Figure 5A1). To avoid confounding effects related to the possibility that not every cell responding to airpuff stimulation is targeted by transfected N-O axons, in this part of the work we only used PO cells where both AP-responses and N-O-driven suppression were confirmed (see METHODS for a detailed description of inclusion and exclusion criteria). Furthermore, we delivered the airpuff 1 second after starting optogenetic stimulation to match the time frame of the strongest suppression of spontaneous activity in PO (Figure 4C1).

**Figure 5.**
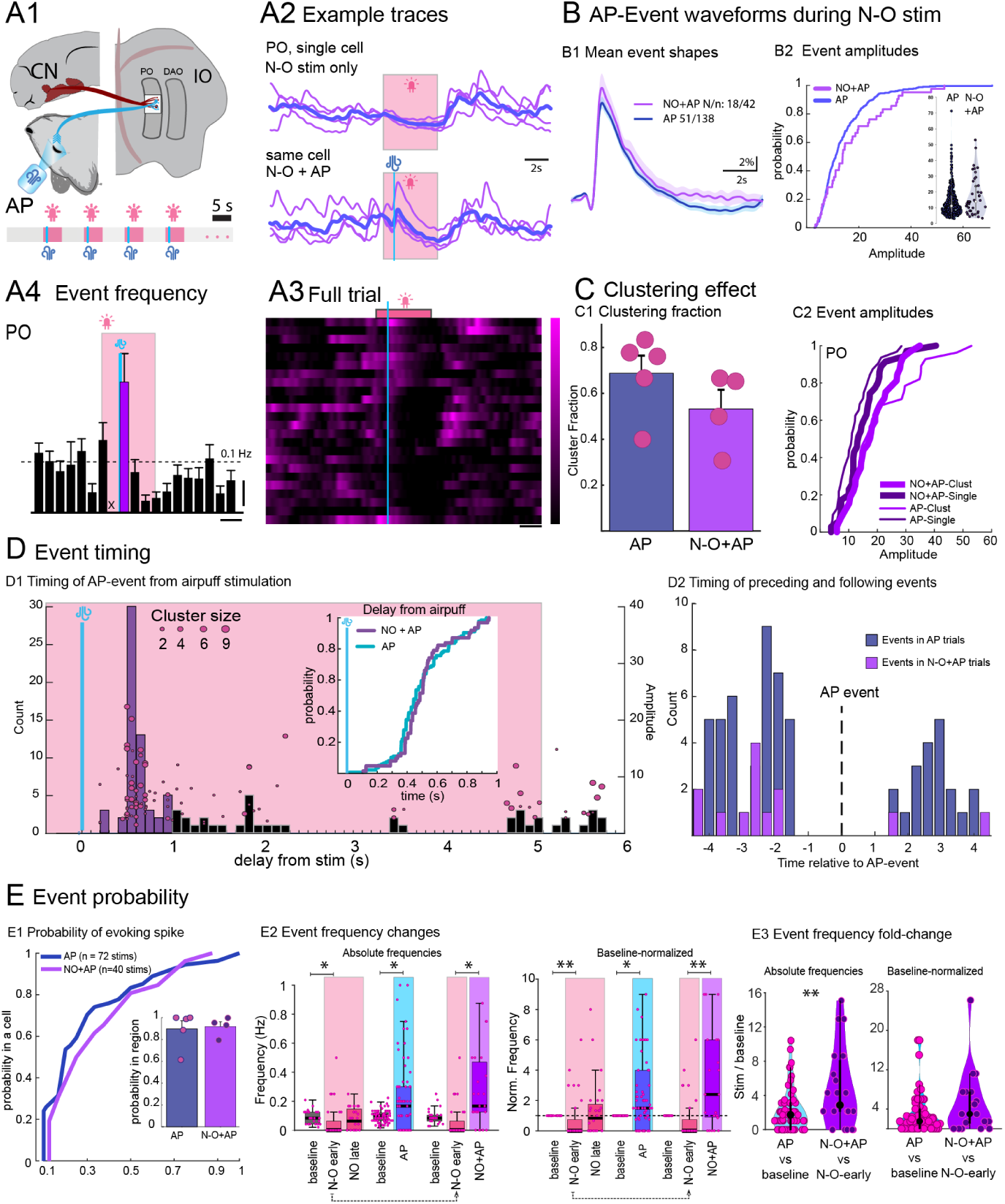
Nucleo-olivary stimulation does not suppress airpuff-evoked spikes. (A) Periocular stimulation evokes IO spikes during N-O stimulation. (A1) Schematic of the experiment. (A2) Example calcium traces from a single cell. N-O stimulation suppresses spontaneous spiking (top), but not AP-evoked spikes (bottom). (A3) Raster plot of the trial (6 cells, 4 stimulations); sorted by event fluorescence intensity. (A4) Stimulus-triggered histogram of event frequencies. (B) N-O stimulation does not decrease AP-evoked event size. (B1) Mean event waveforms (±SEM) for AP-evoked events with (purple) and without (blue) N-O stimulation. (B2) Cumulative distributions and violin plots of event amplitudes with and without N-O stimulation. Inset shows violin plots of the same data. (C) Effect of N-O stimulation on AP-event clustering. (C1) AP-event clustering fraction with and without N-O stimulation; markers represent single recordings. (C2) Clustered and single AP-event amplitudes. Data from AP-events without N-O stimulation shown as a reference. (D) Timing of AP-evoked events. (D1) Purple bars denote events occurring within the 1-second window after stimulation; black bars are later spontaneous events. Circles represent a single spike; vertical position indicates event amplitude, and size indicates cluster size. Inset shows event timing compared to AP-events without N-O stimulation. (D2) Event times preceding and following AP-events (dashed line) without (blue) and with (purple) N-O stimulation. (E) N-O stimulation does not decrease the probability of evoking an IO spike with AP stimulation. (E1) Probability distribution for an AP-event in cells that responded at least once. Inset shows probability of evoking at least one spike in a field of view with a single AP is not affected by N-O stimulation; data from recordings with > 3 AP-responding cells. (E2) Summary of spike frequency with N-O alone (pink), AP alone (cyan), and AP combined with N-O (purple), shown in absolute frequencies (left) and normalised to baseline (right) (E3) Relative increase in event frequency with AP stimulation without and with N-O stimulation, shown in absolute frequency ratio (left) and baseline normalised (right). Abbreviations: CN, cerebellar nuclei; IO, inferior olive; AP, airpuff stimulation; PO, principal olive; DAO, dorsal accessory olive; N-O+AP, airpuff delivery during nucleo-olivary stimulation; Clust, clustered spike; Single, single spike; N-O early, early bin of N-O stimulation block. *, p < 0.05; **, p < 0.01; ***, p < 0.001 **Figure 5—video 1.** Effect of AP during N-O in PO.

To our surprise, PO cells still responded to airpuff stimulations with clear events that increased the average event rate above baseline (Figure 5A2-4). Even more, the waveforms AP events un-der N-O stimulation (“NO+AP”) were not smaller than AP-only-evoked events (AP: 16.26 ± 10.18%, NO+AP: 17.52 ± 12.18%, *p*_GLMM_ = *n*.*a*.; see TABLE 1; Figure 5B1-2). As expected based on the notion that the GABAergic nucleo-olivary axons reduce gap-junctional coupling and thus should reduce synchronous activity in the IO, the larger amplitudes of NO+AP-events could not be explained by event clustering (Figure 5C1). In particular, virtually no clustering-driven amplitude modulation of NO + AP events was observed (Figure 5C2).

To ascertain that the events we classify as AP-evoked events during N-O stimulation are indeed generated by the airpuff and not, for example, rebound events driven by the onset of inhibition, we again examined the timing of events in the post-stimulatory period. As shown in Figure 5D1, there was a clear sharp peak in the stimulation-aligned event timing histogram and the distribution of delays from the sensory stimulation were indistinguishable between AP+NO and AP events (AP: 0.50 ± 0.19*s*, NO+AP: 0.48 ± 0.17*s*, *p*_GLMM_ = 0.48; see TABLE 17; inset in Figure 5D1). Furthermore, the intervals between the AP+NO-events and those preceding and following them were similar to what was observed with AP-only events (Figure 5D2). As discussed earlier (related to Figure 2), relatively invariant spike timing would be expected to be one of the signatures of intrinsically-generated rebound responses (REF). However, it would be hard to explain how rebound events would be consistently generated after 1 second of inhibition (but not after shorter or longer periods). Thus, we cannot avoid the conclusion that the activity of the N-O pathway does not significantly alter the characteristics of sensory-evoked events.

**Table 17.** Delay of AP and N-O+AP events in PO neurons (s). GLMM analysis.

| Group | animalNum | recNum | roiNum | eventNum | Mean | Median | STD | SEM | pValue | hValue |
| --- | --- | --- | --- | --- | --- | --- | --- | --- | --- | --- |
| AP | 5 | 5 | 51 | 138 | 0.50131 | 0.45983 | 0.18709 | 0.015926 |  |  |
| N-O + AP | 5 | 5 | 18 | 42 | 0.48199 | 0.44657 | 0.17181 | 0.02651 | 0.48293 | 0 |

**Notes:** Figure5 D1

Even more, N-O activation did not decrease the likelihood of a single stimulation evoking a response in a cell (mean event probability in AP+NO, 0.28 ± 0.26 (n=40 stimulations); in AP, 0.23 ± 0.25 (n=72 stimulations); *p*_Mann-Whitney_ = 0.918 calculated for recordings where at least 3 cells were responding at least once; see TABLEs 18 and 19 for details; Figure 5E1). The probability of evoking at least one event per stimulation in the visible local region was also unchanged (N-O+AP, 0.83 ±0.1; AP, 0.87 ± 0.07; *p*_Mann-Whitney_ = 0.63; Figure 5E1, inset).

**Table 18.** Event probability in a PO cell. AP vs. N-O+AP. Mann-Whitney U test.

| Group | roiNum | Mean | Median | STD | SEM | pValue | hValue |
| --- | --- | --- | --- | --- | --- | --- | --- |
| AP | 68 | 0.22721 | 0.16667 | 0.25054 | 0.030382 |  |  |
| N-O+AP | 32 | 0.27734 | 0.25 | 0.26066 | 0.046079 | 0.2865 | 0 |

**Notes:** Figure 5 E1

**Table 19.** Event probability in PO subnuclei. AP vs. N-O+AP. Mann-Whitney U test.

| Group | recNum | Mean | Median | STD | SEM | pValue | hValue |
| --- | --- | --- | --- | --- | --- | --- | --- |
| AP | 5 | 0.89774 | 0.9883 | 0.16556 | 0.074042 |  |  |
| N-O+AP | 4 | 0.91907 | 0.94227 | 0.087889 | 0.043944 | 0.66667 | 0 |

**Notes:** Figure5 E1 inset

Puzzled with this lack of suppression, we finally reflected on the fact that the AP+NO-driven event rate should be compared not with the overall baseline activity, but with that expected to be seen during activation of the N-O pathway alone (see summaries of N-O, AP and N-O+AP trials in Figure 5E2 and Tables 20, 21). Indeed, when the AP-evoked event rates were normalized with respect to either pre-stimulation window (for AP-only events, n=69) or to the 1-s bin corresponding to AP stimulation during N-O-only trials (for AP+NO-events, n=23), we found no decrease in event rates (AP vs. baseline [absolute]: 2.365 ± 2.59, N-O+AP vs. N-O-early [absolute]: 4.71 ± 4.49, *p*_Bootstrap_ = 0.006; AP vs. baseline [baseline-norm]: 3.05 ± 4.21, N-O+AP vs. N-O-early [baselinenorm]: 4.86 ± 6, *p*_Bootstrap_ = 0.726; Figure 5E3; see TABLEs 22 and 23.

**Table 20.** Peristimulus event frequency (s). Bootstrap analysis.

| Group | animalNum | recNum | roiNum | stimNum | Mean | Median | STD | SEM | pValue | hValue |
| --- | --- | --- | --- | --- | --- | --- | --- | --- | --- | --- |
| N-0 baseline | 8 | 10 | 28 | 144 | 0.0875 | 0.0833 | 0.0455 | 0.0086 |  |  |
| N-0 early | 8 | 10 | 28 | 144 | 0.0580 | 0 | 0.1202 | 0.0227 | 0.0140 | 1 |
| APbaseline | 5 | 5 | 69 | 631 | 0.0960 | 0.1 | 0.0431 | 0.0052 |  |  |
| APfirstStim | 5 | 5 | 69 | 631 | 0.2268 | 0.1667 | 0.2487 | 0.0299 | 0.0280 | 1 |
| N-0+AP baseline | 5 | 5 | 23 | 162 | 0.0963 | 0.0833 | 0.0544 | 0.0113 |  |  |
| N-0+AP | 5 | 5 | 23 | 162 | 0.2736 | 0.1667 | 0.2605 | 0.0543 | 0.1520 | 0 |
| N-0 early | 8 | 10 | 28 | 144 | 0.0580 | 0 | 0.1202 | 0.0227 |  |  |
| N-0+AP | 5 | 5 | 23 | 162 | 0.2736 | 0.1667 | 0.2605 | 0.0543 | 0.0040 | 1 |

**Notes:**

**Table 21.** Peristimulus event frequency [Normalized to baseline]. Bootstrap analysis.

| Group | animalNum | recNum | roiNum | stimNum | Mean | Median | STD | SEM | pValue | hValue |
| --- | --- | --- | --- | --- | --- | --- | --- | --- | --- | --- |
| N-0 baseline | 8 | 10 | 28 | 144 | 1 | 1 | 0 | 0 |  |  |
| N-0 early | 8 | 10 | 28 | 144 | 0.80 | 0 | 1.58 | 0.30 | 0.0020 | 1 |
| APbaseline | 5 | 5 | 69 | 631 | 1 | 1 | 0 | 0 |  |  |
| APfirstStim | 5 | 5 | 69 | 631 | 3.05 | 1.50 | 4.21 | 0.51 | 0.0360 | 1 |
| N-0+AP baseline | 5 | 5 | 23 | 162 | 1 | 1 | 0 | 0 |  |  |
| N-0+AP | 5 | 5 | 23 | 162 | 3.90 | 2.40 | 4.83 | 1.01 | 0.1100 | 0 |
| N-0 early | 8 | 10 | 28 | 144 | 0.0580 | 0 | 0.1202 | 0.0227 |  |  |
| N-0+AP | 5 | 5 | 23 | 162 | 3.90 | 2.40 | 4.83 | 1.01 | 0.0060 | 1 |

**Notes:**

**Table 22.** Peristimulus event frequency fold-change. Bootstrap analysis.

| Group | animalNum | recNum | roiNum | stimNum | Mean | Median | STD | SEM | pValue | hValue |
| --- | --- | --- | --- | --- | --- | --- | --- | --- | --- | --- |
| AP vs. baseline | 5 | 5 | 69 | 631 | 2.3623 | 1.7358 | 2.5904 | 0.3118 |  |  |
| N-0+AP vs. N-0 early | 5 | 5 | 23 | 162 | 4.7135 | 2.8718 | 4.4879 | 0.9358 | 0.0060 | 1 |

**Notes:** Figure5 E3 Absolute

**Table 23.**
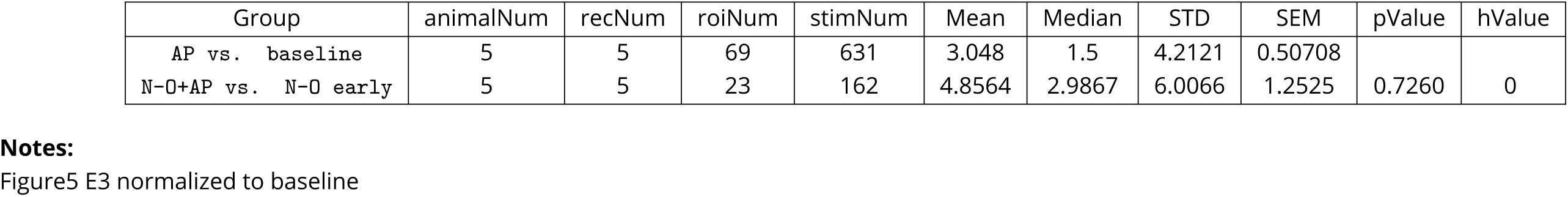
Peristimulus event frequency fold-change [Normalized to baseline]. Bootstrap analysis.

| Group | animalNum | recNum | roiNum | stimNum | Mean | Median | STD | SEM | pValue | hValue |
| --- | --- | --- | --- | --- | --- | --- | --- | --- | --- | --- |
| AP vs. baseline | 5 | 5 | 69 | 631 | 3.048 | 1.5 | 4.2121 | 0.50708 |  |  |
| N-0+AP vs. N-0 early | 5 | 5 | 23 | 162 | 4.8564 | 2.9867 | 6.0066 | 1.2525 | 0.7260 | 0 |

**Notes:** Figure5 E3 normalized to baseline

## Discussion

Our study presents the first multicellular recordings from the inferior olive (IO) in a living animal, offering valuable insights into the origins of cerebellar complex spike dynamics and the effects of nucleo-olivary (N-O) feedback into the IO. Our results bring light to two central issues in the field of olivo-cerebellar systems. First, we provide information on the determination of the spike sizes of the cerebellar complex (***Najafi et al. (2014b)***). Second, we show that the effect of cerebellar nucleo-olivary activation can be context-dependent.

### Differences between olivary subnuclei reinforce modulary specialization of the cerebellar function

The cerebellum has historically been considered to have a homogeneous microzonal structure and function (***Middleton and Strick (1998)***), allowing researchers to generalize findings from specific regions to the entire olivo-cerebellar system. However, reports of differences in function (***Wadiche and Jahr (2005); Wang et al. (2011)***) have been consolidated by Zhou et al.’s systematic study of Purkinje neuron activity across the cerebellar cortex in awake animals (***Zhou et al. (2014)***). This foundational work established clear differences between Purkinje neurons (PNs) in zebrin-positive (Z+) and zebrin-negative (Z-) microzones, with Z+ PNs displaying lower simple spike (SS) and complex spike (CS) frequencies compared to Z-PNs. The lower SS rate was attributed to intrinsic factors of PNs, while the reduced frequency of CSs was speculated to result from increased inhibition of the IO via the nucleo-olivary (N-O) pathway targeting the same microzone.

Our results from the IO regions targeting Z+ (principal olive, PO) and Z- (dorsal accessory olive, DAO) zones align with these findings. Neurons in the PO consistently exhibited lower event rates (Figure 1C and Figure 2A, C-D). The somewhat weaker inhibition of spontaneous activity in the PO compared to DAO after activation of the N-O pathway (Figure 4C) could reflect greater tonic inhibition in the PO, although further experiments, such as direct observations of N-O axon activity levels in PO and DAO, are needed to confirm this.

Zhou and colleagues also noted that Z+ CSs had broader waveforms than Z-CSs (***Zhou et al. (2014)***), a phenomenon that we observed as larger event amplitudes in the PO compared to the DAO. Although interpreting the amplitude of the calcium signal as a proxy of the duration of the IO spike requires caution, we found that the event amplitudes in PO cover a greater range than in DAO (Figure 2B). Further examination revealed that co-activation (“clustering”) with neighboring cells increased event amplitude (Figure 2E1). Noting the higher amplitudes of PO events, we hypothesized that neurons in PO could exhibit stronger clustering, possibly due to differences in electrical connectivity within the IO (***Hoge et al. (2011); Vrieler et al. (2019)***). Indeed, the clusters of events involved a greater proportion of neurons in the PO than in the DAO (Figure 2E2) and fewer events occurred in isolation (Figure 2E3). Thus, a bias toward larger coactivation groups in the Z+ targeting region could explain the broader complex spikes observed by Zhou et al.

Despite searching, we found no reports that examined coactivation levels between Z+ and Z-microzones in cerebellar complex spike recordings. However, our findings are consistent with studies in anesthetized rats showing that the duration of the complex spike is positively correlated with local levels of coactivation (***Lang et al. (2014)***).

### Enhanced coactivation of PO neurons with sensory stimulation may explain larger sensory-evoked CSs

Determining the exact mechanism behind the amplitude-enhancing effect of spontaneous event coactivation remains beyond the scope of this study. However, we observed a similar trend in sensory-evoked events: airpuff (AP)-evoked events were larger and exhibited higher levels of coactivation (Figure 3B, C). Moreover, the increase in event amplitude corresponded to differences in coactivation, such that the clustered AP-evoked events were indistinguishable in amplitude from the clustered spontaneous events (Figure 3C2). These findings suggest that sensory stimulation enhances coactivation among PO neurons, which may contribute to the generation of larger amplitude complex spikes, providing further insight into how sensory inputs modulate cerebellar output.

Interestingly, during N-O stimulation, the relatively few spontaneous events observed in the PO and DAO were not smaller than other spontaneous events recorded in the same cells, even though they occurred with less coactivation than expected based on event rates, supporting the notion that N-O axons affect electrical coupling (Figure 4D ***Lefler et al. (2014)***). One possible explanation is that the events were initiated from a lowered baseline calcium level due to N-O activation (Figure 4C2), which might expand the available dynamic range of GCaMP6, allowing for greater fluorescence fluctuations. However, our results align with a recent report showing that activation of the nucleoolivary pathway did not reduce the number of spikelets in complex spikes evoked by periorbital stimulation in decerebrate ferrets (***Öhman et al. (2024)***).

Overall, our GLMM analysis rejected the hypothesis that event amplitude variability would be mainly due to random effects such as variable expression of GCaMP6 or intrinsic properties of individual neurons. Instead, our data suggest that the larger complex spikes observed in individual Purkinje neurons in response to sensory stimulation (***Najafi et al. (2014a); De Gruijl et al. (2014); Ju et al. (2019)***) partly reflect the broader co-activation of olivary neurons compared to spontaneous events. Considering that Purkinje neuron excitability can influence the final form of the complex spike (***Zang and De Schutter (2019)***), the complex spike waveform emerges as a signal that can be subtly modified at multiple stages of cerebellar computation.

### Influence of nucleo-olivary feedback on inferior olive excitability

One of the central concepts in cerebellar theories is the cerebellum’s ability to modulate its own complex spike activity (***Kistler and De Zeeuw (2002); Bengtsson and Hesslow (2006); Rasmussen and Hesslow (2014); Herreros and Verschure (2013); Tokuda et al. (2013); Chaumont et al. (2013)***). While di- or polysynaptic signaling loops (where CN projections influence regions that, in turn, send afferents back to the inferior olive (IO)) likely play roles over long time scales (***van Hoogstraten et al. (2024); Ruigrok et al. (2023)***), the direct feedback via the nucleo-olivary (N-O) pathway has garnered the most attention. Until recently, the N-O pathway was believed to be exclusively GABAergic. However, detailed anatomical and physiological investigations have now established that a portion of the cerebellar nuclear projection to the medial accessory olive (MAO) is glutamatergic (***Judd et al. (2021); Wang et al. (2023)***).

Despite this added capacity for direct excitation, modulation of climbing fiber activity by the N-O pathway is generally considered to occur either through the uncoupling of dendritic gap junctions between IO neurons (***Llinas (1974); Lefler et al. (2014); Marshall and Lang (2009)***) or by suppressing the excitability of IO neurons via conventional inhibitory mechanisms (***Andersson et al. (1988)***). This view aligns with evidence showing that the excitability of the IO decreases during preparation for a conditioned response (***Sears and Steinmetz (1991)***). However, this suppression has not been directly demonstrated to result from activity in N-O axons. Furthermore, past experiments aiming to reveal suppression of IO excitability related to the N-O pathway have not been entirely straightforward to interpret (***Ruigrok and Voogd (1995); Apps and Lee (2002)***).

To determine whether activation of the nucleo-olivary (N-O) pathway can lead to changes in inferior olive (IO) spiking consistent with excitability suppression via classic GABA-receptor-mediated hyperpolarization, we examined the rates of spontaneous and sensory-evoked IO events during optogenetic stimulation of the N-O pathway. To our surprise, while N-O stimulation robustly suppressed spontaneous activity (Figure 4),sensory-evoked events were unaffected (Figure 5). This finding was particularly striking because we limited our analysis to recordings in which we could confirm both the presence of events evoked by periocular airpuff (AP) as well as clear suppression of spontaneous activity caused by N-O stimulation. Furthermore, we excluded all trials and cells in which N-O stimulation led to direct excitation (see Methods for detailed inclusion and exclusion criteria). Despite this further restriction of our data set, airpuff-evoked event rates remained at least as high under N-O activation as under baseline conditions (Figure 5E).

### Refractory period as a central factor limiting IO activity

Our findings present a puzzling contradiction: while activation of the nucleo-olivary (N-O) pathway robustly suppresses spontaneous activity in the inferior olive (IO) to near silence. This is unexpected because it is well established by us and others(***Lefler et al. (2014); Loyola et al. (2023); Bazzigaluppi et al. (2012b); Best and Regehr (2009)***) that action potentials in N-O terminals lead to hyperpolarizing postsynaptic responses in IO neurons, and activation of N-O axons can lead to effective suppression of airpuff-evoked spikes in awake animals (***Kim et al. (2020)***). If N-O activation hyperpolarizes IO neurons, we would expect both spontaneous and sensory-evoked activity to be suppressed.

One possible explanation is that our sensory stimulation was unnaturally strong relative to the inhibitory capacity of the N-O pathway, potentially overriding the inhibition. However, the probability that a given cell would respond to an airpuff in our experiments was relatively low (0.22 ± 0.25) compared to the values reported in previous studies (0.2–0.6; ***Najafi et al. (2014b); Yang and Lisberger (2017); Ju et al. (2019); Lin et al. (2024)***). This suggests that our sensory stimulation was not excessively powerful and is unlikely to have overridden the inhibitory effects of N-O activation.

Another potential factor could be the time course of N-O-mediated inhibition. Although our optogenetic stimulation parameters were based on previous *in vitro* work demonstrating that N-O synapses can maintain tonic responses during 20 Hz activation (***Lefler et al. (2014)***), it is possible that in intact circuitry, the effective suppression is more transient. However, we timed airpuff stimulation to coincide with the period of strongest suppression of spontaneous spiking (1–2 seconds into the stimulation period), making transient inhibition an unlikely explanation.

We propose that the key to reconciling these effects lies in the refractory period of IO neurons. Analysis of the intervals between spontaneous and airpuff-evoked events (Figure 5D2) revealed that under our experimental conditions, no events occurred less than 1.5 seconds apart, regardless of whether they were spontaneous or evoked. This substantial refractory period suggests that IO neurons are unlikely to generate spikes shortly after a preceding event, whether spontaneous or sensory-evoked. While this refractory period is bound to be shorter in awake animals, it indicates that refractoriness severely limits the possibility of spike generation in our preparation.

Inhibition of spontaneous events during continuous N-O activation reduces the likelihood that IO neurons are in a refractory state when sensory inputs are presented. This in turn can increase the probability that afferent inputs will initiate a spike with minimal delay. Notably, after N-O activity ends, the increased availability of IO neurons recovered from refractoriness might contribute to the reported “rebound” activity (***Kim et al. (2020)***).

This mechanism provides a plausible explanation for our observation that sensory-evoked events remain unaffected during N-O activation despite suppression of spontaneous activity. It also aligns with reports that the magnitude of complex spikes increases with longer intervals between complex spikes (e.g., ***Maruta et al. (2007)***), possibly due to a larger pool of IO neurons available for coactivation after a longer interval. As our data indicate that the amplitude of the IO event increases with the number of coactivated neurons (Figure 2 E), the larger sizes of the late complex spikes may be a reflection of a larger pool of available IO neurons to respond.

In summary, the dependence of the excitability of the IO neuron on the preceding event interval suggests that activation of the GABAergic N-O pathway could enhance the transmission of sensory information to the IO by reducing spontaneous activity and refractoriness. By diminishing background activity, this mechanism would allow sensory inputs to evoke IO spikes with precise timing, unimpeded by prior spontaneous spiking. Although we must consider the slower time scales inherent in our preparation, these insights are valuable to understand IO activity during behavior and the generation of complex spikes.

## Methodological considerations

The inferior olive (IO) is located in the ventral medulla, surrounded by several life-critical nuclei, making recordings in intact animals extremely challenging. Some reports of IO activity in living animals have used a caudal approach through the *foramen magnum* (***Chorev et al. (2007)***), but even though appropriate for single-electrode recordings, this method is incompatible with singlecell-resolution fluorescence imaging.

Here, inspired by earlier works employing single-electrode recordings (***Bazzigaluppi et al. (2012a); Khosrovani et al. (2007)***) we leveraged the fact that the medullary region housing the mouse IO protrudes beyond the ventral skull’s confines. By slightly enlarging the *foramen magnum* during a ventral approach surgery (detailed in ***Guo et al. (2021)***), portions of the IO become directly accessible and neuronal activity can be optically monitored without breaking the *dura mater*. The surgery is obviously non-reversible and the animal cannot be allowed to wake from deep anesthesia; furthermore, the spontaneous activity is significantly lower (∼0.1 Hz) than in awake animals (commonly reported as 1–2 Hz; ***Titley et al. (2019)***). Despite these shortcomings, the simultaneous access to two distinct subnuclei of the IO, the non-compromised system-level connectivity, and the possibility of targeted optogenetic manipulation make this approach a powerful tool for addressing central questions about olivo-cerebellar function.

Importantly, the intrinsically slow event rate of the IO makes it suitable for fluorescence-based activity monitoring within the olivo-cerebellar system, where many neurons exhibit spontaneous high-frequency firing (***Heck et al. (2013)***). Prioritizing sensitivity and dynamic range over rapid rise times, we chose GCaMP6s due to its excellent dynamic range (***Zhang and Looger (2024)***), which we previously demonstrated can report changes in the duration of the action potential of IO (***Dorgans et al. (2022)***).

However, interpreting fluorescence signals relative to underlying voltage changes is challenging, particularly in IO neurons with unusual calcium dynamics (***Benardo and Foster (1986); Llinas and Yarom (1981); Schweighofer et al. (1999); Bazzigaluppi and De Jeu (2016)***). Furthermore, the absence of optical sectioning in the whole-field imaging method can lead to confounding artifacts in densely labeled structures such as the IO’s tortuous neuropil. As advanced technical solutions such as structured illumination (***Li et al. (2020)***) or two-photon microscopy (***Zong et al. (2022); Nardin et al. (2024)***) were beyond our reach, we made use of the computational method CNMF-E (Constrained Non-negative Matrix Factorization for Endoscopic imaging; ***Zhou et al. (2018)***) to extract signals assigned to individual cell somata. This method assumes event sparsity in both time and space, as well as non-negativity; violating these assumptions can lead to incorrect results (***Vanwalleghem et al. (2021)***). Although IO activity is temporally sparse, spatial sparsity may be compromised due to intrinsic synchronization via gap junctions and neuropil activity. In the absence of ground-truth data to validate the CNMF-E results, we carefully examined each recording and excluded any suspicious components. Recognizing this element of subjectivity, we will make the imaging data publicly accessible at [zenodo repository address] to ensure full transparency.

## Conclusions and future directions

Our study presents a novel methodology that bridges the gap between *in vitro* and *in vivo* studies, allowing multicellular recordings from the inferior olive (IO) in living animals. This approach could potentially be adapted to investigate other deep brainstem structures that have historically been challenging to study.

Our methodology serves as an “upgraded” version of *in vitro* experimentation, partly retaining the accessibility and resolution of slice preparations while also incorporating the intact neural circuitry and physiological conditions of a living organism. Although this approach cannot replace studies conducted in awake, behaving animals, it complements them by providing detailed cellular and network-level insights akin to those derived from decades of *in vitro* research that laid the foundation for current systems neuroscience.

In particular, experimentation within the IO presents an opportunity for examining the organization and function of the olivocerebellar micromodules, which are presumed to form closed loops (***Apps et al. (2018); De Zeeuw (2021); Payne et al. (2024)***). Among the elements that form this loop, the IO represents the most compact and uniform structure, with well-defined and restricted projections. Therefore, in contrast to other components of the olivo-cerebellar circuits with numerous cell types and extensive local interneuronal connectivity, the relative simplicity and homogeneity of the IO make it an ideal target for monitoring interactions within and between cerebellar micromodules.

Nevertheless, future studies should aim to achieve multicellular records of IO activity in awake, behaving animals to reveal how olivary dynamics support behavioral coordination under natural physiological conditions.

## Methods and Materials

### Animal ethics statement

Male C57BL/6J mice aged postnatal (P)35-45 were used for this study. All animal experiments were conducted in accordance with the 2006 guidelines for Proper Conduct of Animal Experiments of the Science Council of Japan and were approved by the Committee for Care and Use of Animals at the Okinawa Institute of Science and Technology, an AAALAC-accredited facility under protocols 2021-005 and 2024-021.

### Stereotaxic virus vector injection

Following established protocols (***Guo et al. (2021); Dorgans et al. (2022)***), stereotaxic injections were performed to express GCaMP6s in the IO neurons and ChrimsonR-tdTomato in CN neurons. Postnatal (P)35-45 male C57Bl/6J mice were anesthetized with 5% isoflurane and mounted in a stereotaxic frame (Motorized Stereotaxic based on Kopf Model 900. Neurostar, Germany). Anesthesia was maintained with 1.8–2.4% isoflurane delivered via a nose cone. The scalp hair was removed using a shaver and Veet depilatory cream (Veet sensitive skin. Veet, Canada). The exposed skin was locally anesthetized with Xylocaine gel (Xylocaine Jelly 2 %. Aspen, Japan) and disinfected with 70% ethanol. A midline incision was made along the exposed scalp to reveal the skull.

To ensure precise targeting of the IO, located at the most ventral part of the medulla, the skull was carefully leveled by minimizing the difference between bregma and lambda on the z-axis to within 0.05 mm. Small craniotomies ( 1 mm in diameter) were then created using a hand-held drill (Surgic XT Plus drill, NSK, Japan) to access the IO and contralateral CN.

Viral vectors were diluted in sterile saline and loaded into a customized quartz glass pipette (Q114-53-10NP. Sutter Instrument, CA, USA) pulled with laser puller (P-2000. Sutter Instrument, CA, USA). For IO injections, a mixture of AAV9.TRE.GCaMP6s.WPRE and AAV9.5HTr2b(3.7)-tTA.WPRE was injected. The pipette was inserted at a rate of 0.2 mm/s into two coordinates (relative to bregma) to ensure broader coverage of the IO along the anterior-posterior dimension. The first injection was made at AP -6.2 mm, ML ±0.5 mm, and DV 6.7 mm (posterior IO), and the second at AP -6.2 mm, ML ±0.5 mm, and DV 6.6 mm (anterior IO). At each site, 150 nl of viral solution was delivered using a nanojector (Neurostar, Germany) at a rate of 30 nl/min. For the contralateral CN injection, a new quartz glass pipette filled with AAV9.Syn.ChrimsonR-tdTomato.WPRE.bGH was used. Two injections were delivered at the following coordinates to cover most interposed and lateral CN: AP -6.3 mm, ML ±1.75 mm, DV 3.6 mm; and AP -6.1 mm, ML ±1.8 mm, DV 3.5 mm. Similar to IO injection, 150 nl of viral solution was delivered at each site at a rate of 30 nl/min.

In total, 600 nl of viral solution was injected per animal. After injections, the scalp was sutured and disinfected with 70% ethanol. For postoperative pain management, 0.05 mg / ml of Rimadyl (Zoetis, New Jersey, US) was administered subcutaneously at a dose of 5 mg/kg.

### In vivo calcium imaging of IO neurons

14 to 21 days after viral injection, the medulla was accessed from the ventral side of anesthetized animals following a previously described protocol (***Guo et al. (2021)***). Animals were anesthetized using the same protocol as for the viral injection, then mounted in a stereotaxic frame in a supine position, with the ventral side facing up. Neck hair was removed using a shaver and Veet depilatory cream, and the neck skin was incised. The salivary glands were freed from connective tissue and reflected laterally to expose the trachea.

To prevent discomfort from potential isoflurane insufficiency during the tracheotomy, animals received an intraperitoneal injection of 7.5 mg/ml Ketamine (Daiichi Sankyo, Japan), administered twice at 5-minute intervals to achieve a total dose of 50 mg/kg. The trachea was secured to the chest skin and intubated for continued isoflurane delivery. The esophagus and muscles overlying the foramen magnum were carefully removed. The ventral arches of the atlas (the first cervical vertebra) were cut to remove the anterior tubercle of the atlas, providing a clear view of the foramen magnum. Finally, the foramen magnum was enlarged by removing part of the occipital bone to access the superficial parts of the principal olive (PO) and dorsal accessory olive (DAO) for imaging. 15 minutes prior to calcium imaging, animals were administered alpha-chloralose intraperitoneally at a dose of 114 mg/kg. Alpha-chloralose was chosen over isoflurane because it does not interfere with gap junctions, which are essential for proper neuronal signaling in the inferior olive. However, since alpha-chloralose has a weak analgesic effect, isoflurane was delivered between recordings to ensure the animals did not experience pain. A mini-microscope (nVoke2. Inscopix, CA, USA), coupled with a gradient-index (GRIN) lens (1 mm diameter, 9 mm length, 0.5 NA in water), was used to image the superficial parts of the inferior olive without penetrating the dura mater. GCaMP6s signals were excited with a blue LED (455 ± 8 nm) at a power of 1.2–1.8 mW. For optogenetic stimulation of ChrimsonR, a red LED (620 ± 30 nm) was used at a power of 1 mW. Power was measured at the front surface of the objective.

Periocular airpuff stimulation was delivered via a plastic tube with a 1mm diameter tip, positioned approximately 1.5 cm from the contralateral eye, connected to a pressure injector (MPPI-3, ASI, OR, USA) set at 120 kPa for 100 ms. Calcium signals were recorded at a sampling frequency of 20 Hz.

### Calcium imaging data preprocessing

Calcium imaging movies were first spatially filtered, then motion corrected, and subsequently exported to TIFF files using the MATLAB API of Inscopix Data Processing Software. Spatial filtering was applied to each frame (**M**_*f*_ ) individually using a bandpass filter designed to enhance feature visibility and reduce noise. The filter was implemented by subtracting a Gaussian-blurred version of the frame with a low cutoff frequency (*σ*_low_ = 0.5 cycles per pixel), which retains higher spatial frequencies, from a Gaussian-blurred version with a high cutoff frequency (*σ*_high_ = 0.005 cycles per pixel), which retains lower spatial frequencies. This process can be mathematically represented as:

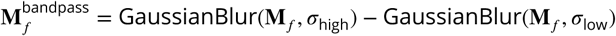

The spatial filtering also enhanced the accuracy of the subsequent motion correction by emphasizing relevant cellular features. Motion correction was conducted using an image registration method described in ***Thevenaz et al. (1998)*** [REF:Thevenaz1998]. The first frame of the movie was selected as the global reference frame. Each subsequent frame was aligned to this reference frame to address any potential drift across the entire movie sequence. The motion correction was further constrained with a maximum translation limit of 20 pixels in any direction, ensuring realistic and appropriate corrections.

After being converted to TIFF files, data analysis was performed using MATLAB (R2024a. Math-Works, Massachusetts, US). The motion-corrected movies were processed by the MATLAB package of Constrained Nonnegative Matrix Factorization for microEndoscopic data (CNMF_E; https://github.com/zhoupc/CNMF_E) to denoise the recording and extract individual neural activity (***Zhou et al. (2018)***). In brief, CNMF-E models the calcium imaging data *Y* as a combination of spatial, temporal, background, and noise components:

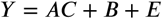

Discrete spatial components (*A*), coupled with corresponding discrete temporal activity (*C*), were identified to represent ROIs that correspond to neuron somas. To denoise the data, CNMF_E estimated the gradual background changes using data from regions outside the identified ROIs and the neuropil signal by averaging the activity from pixels near neuron somas. Both the slowvarying background and neuropil contamination (*B*) were subtracted from the raw fluorescence data.

After obtaining the denoised fluorescence traces, CNMF-E applied a deconvolution step to infer spike events by modeling calcium transients using an autoregressive process. During this process, the residual noise (*E*), which represents the unexplained variability in the data, is minimized. The deconvoluted signal reflects the likely timing and magnitude of neural spikes.

For both the denoised and deconvoluted data, Δ*F* ∕*F* was calculated by normalizing the fluorescence signal to the estimated baseline fluorescence (*F*_0_), which represents the neuron’s resting fluorescence during periods of minimal neural activity.

Both the denoised and deconvoluted temporal components were used for analysis. The deconvoluted component contains reshaped calcium transients modeled by an autoregressive process, making it easier to isolate and identify calcium events. The time information of these calcium transients was then used to locate events in the denoised data, which retains more of the original structure of the data, including the slow-varying baseline fluorescence and other non-modeled transients.

### Calcium imaging data processing and calculation

The denoised data was processed separately using a 1 Hz low-pass filter to smooth the traces and a 4 Hz high-pass filter to extract the noise signal. The noise signal from the high-pass filter was used to normalize calcium event amplitudes across neurons and recordings. Various properties were then measured based on the processed traces.

The rise point of each calcium transient was determined as the last point within a 1-second window prior to the peak where the calcium signal was either decreasing or remaining constant, followed by an increase. The amplitude of each transient was calculated as the difference between the calcium signal values at the peak and the rise point. To standardize the measurements, the amplitude was normalized to the standard deviation (STD) of the high-pass filtered data. For the full-width at half maximum (FWHM) calculation, the raw transient amplitude was used to identify the half-maximum value. The points on the rising and decaying phases of the transient where the signal crossed half of the maximum amplitude were determined. To improve accuracy, interpolation between data points was performed using the shape-preserving piecewise cubic interpolation method, with query points spaced at 5 ms intervals. The time difference between the interpolated half-maximum points on the rise and decay phases was reported as the FWHM of the transient.

### Categorization of ROIs and calcium events

The central artery, running along the midline of the medulla on the ventral surface, was used as the primary landmark to separate the left and right hemispheres of the IO. In each hemisphere, additional blood vessels running parallel to the central artery provide a natural anatomical reference, with the PO located medially and the DAO situated laterally.

Calcium events were categorized according to their temporal relationship with various stimulation protocols. Events that occurred before any stimulation or more than 1 second after stimulation were classified as spontaneous events (spon). Events triggered by airpuff stimulation, occurring within 1 second of the stimulus, were grouped as airpuff-triggered events (AP-Evoked). Events activated by optogenetic stimulation (OG) of the N-O pathway within 1 second of the onset of OG stimulation were classified as events triggered by N-O (N-O evoked). Events that appeared during OG stimulation, but with a delay of more than 1 second, were classified as spontaneous events during OG stimulation (sponDuring N-O). Events that occurred immediately after the termination of OG stimulation, within 1 second, were categorized as rebound events after N-O (N-O rebound). Finally, events triggered by airpuff stimulation during ongoing optogenetic stimulation, within 1 second of airpuff onset, were classified as OGAP-triggered events (N-O+AP).

To maintain data quality and ensure the inclusion of only healthy neuron responses, ROIs with a low frequency of spontaneous events (spon) (≤ 0.05*Hz*) were excluded from further analysis. Additionally, calcium events detected in the denoised data were further evaluated based on strict criteria, including a minimum transient rising slope of 3 and a peak-to-noise ratio of at least 3. For some events, where the calcium signal did not decay below half of the maximum amplitude before another rise occurred, FWHM could not be determined. These events were excluded from the FWHM analysis.

### Clustered event definition

We loosely follow the concepts described in ***Lang et al. (1996)*** for synchrony definitions. Viral labeling will always lead to variable fraction of neurons recorded in a given field of view preventing accurate estimates of active cell count. Furthermore, the slow temporal resolution inherent to miniscope-based calcium imaging renders any claims of millisecond-level synchronization meaningless. Instead, we demarcate individual events as “clustered” if a spike with peak time within 50 ms (one frame of recording) was found in at least one other cell. Clustering metric was obtained by pooling event peak times from all cells in the field of view, and clustering the times using the dbscan (***ESTER (1996)***) algorithm, with epsilon = 1 frame and a minimum number of events in a cluster = 2. Clustering fraction was given as the proportion of events in a given recording that were classified as clustered. Relative clustering was calculated as the ratio between observed clustering and expected clustering *S*, the latter calculated based on mean event rates and available cell counts, and assuming event independence, in a given field of view:

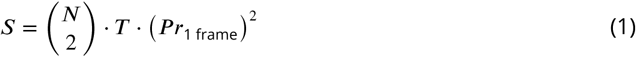

where 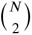 is the number of combinations of cell pairs from *N* cells, *T* is the length of the recording time period in frames, and *P r*_1 frame_ is the average probability of a spike occurring in a single frame interval.

Note that for events evoked with a stimulation, the available cell count only included cells that responded to the stimulation in that given trial.

### Peri-stimulatory Calcium Event Frequency Analysis

To dissect the temporal dynamics of neuronal responses to various stimuli, events were organized into time bins, both standardized to one second and tailored to specific experimental conditions. The one-second bins provided a uniform framework for temporal analysis across all setups, while the customized bins were strategically grouped to enhance the comparison of key experimental phases. This approach allowed for a focused examination of modulation of activity at selected time frames during and after stimulation. Events were aligned with the onset of stimulation for both the one-second and customized bin methods, ensuring a uniform starting point for analysis across all experimental setups. The period from -6 to 0 seconds before the stimulation onset was used as the baseline for both the one-second and customized bin methods to evaluate the effect of stimulation.

Customized bins were designed to elucidate the specific effects of different stimulation protocols. For AP and N-O+AP trials, events within the one-second post-AP stimulation onset were categorized into AP bins and N-O+AP bins, respectively. To isolate the effects of AP in the presence of N-O, the same one-second window used in N-O trials, termed the N-O early bin, served as a comparative baseline. This baseline adjustment ensures that any modulation from N-O is accounted for when assessing the impact of AP in the N-O+AP bin. Each bin’s activity was then normalized to its respective baseline (AP bins to their pre-stimulation period and N-O+AP bins to the N-O early bin). Comparing these normalized values allows for a direct evaluation of AP’s effects both alone and in combination with N-O, effectively isolating how neuromodulation alters the AP response.

Following the one-second N-O early/N-O+AP bins, the periods in N-O and N-O+AP trials were classified as the N-O late bins, spanning three seconds to capture the sustained effects of N-O. After the end of the stimulus—whether AP, N-O, or N-O in N-O+AP trials—a six-second window was designated as the recovery bin to monitor the reversion of neuronal activity to baseline levels, providing insights into the recovery dynamics post-stimulation.

### Histology

After calcium imaging recordings, animals were transcardiac perfused using 4% paraformaldehyde (EMS, PA, USA). Overdose of pentobarbital or secobarbital was administered via an intraperitoneal injection in advance to prevent pain. The brains were extracted and post-fixed in 4% paraformaldehyde at 4 ^◦^C overnight. 100 *μm* brain coronal slices were prepared with a vibratome (Campden, UK) and mounted on glass slides using Vectashield mounting medium (CA, USA) to preserve fluorescence.

Image acquisition was performed with a Zeiss LSM 880 confocal microscope (Zeiss, Germany) with a 16-bit depth in line-scan mode. For GCaMP6s, excitation was achieved using an Argon laser at 488 nm with emission collected between 490-535 nm, while for ChrimsonR-tdTomato, excitation was performed using a DPSS 561 nm laser with emission collected between 470-655 nm.

A 10x objective (Plan-Apochromat 10x/0.45 M27) was used to obtain an overview of the brainstem and cerebellum, while a 20x objective (Plan-Apochromat 20x/0.8 M27) was utilized for detailed imaging of specific nuclei, such as the IO and CN. To capture larger fields of view and gather information from multiple depths, tiling and z-stacks (4 *μm* z-step and a XY pixel size of 0.83 *μm* were used for 10x objective; 2 *μm* z-step and an XY pixel size of 0.415 *μm* were used for 20x objective) were employed together. The tiled images were first stitched using Zeiss Zen software (PRODUCT), and the maximum intensity projections of the stitched z-stacks were then generated in ImageJ to enhance the visualization of fluorescence signals.

### Statistics

All statistical analyses were done using customized code written for MATLAB. The code used for calculation and plotting is available in an online repository (nvoke-analysis; https://github.com/gblackhorn/nvoke-analysis.git).

Bar plots display the MEAN ± SEM; Violin plots indicate the density of data by the width of “violin” and highlight the MEDIAN and quartiles of the data. Empirical cumulative distribution plots show the full data distribution by plotting the cumulative proportion of data points against their values; Box plots summarize the median, quartiles, and extend whiskers to the most extreme data points within 1.5 times the interquartile range.

The data was collected from multiple ROIs and recordings, resulting in nested, non-paired data with potential correlations within clusters. To account for this structure and address within-cluster correlations, the data was analyzed using both Linear Mixed Models (LMM) and Generalized Linear Mixed Models (GLMM). LMM was used to compare the non-skewed data, such as changes in calcium levels during optogenetic stimulation. In contrast, several calcium event properties exhibited skewed distributions, characterized by asymmetry and a long tail. These variables included event amplitude, rise duration, FWHM, as well as event frequency and intervals. A GLMM with a gamma distribution and log link function was applied to analyze these variables, as it accommodates the non-normal, positively skewed nature of the data and their strictly positive values.

To compare the response variable between two groups, two models were constructed: a full model including the group variable as a fixed effect, and a reduced model excluding this group variable. The group variable represents different experimental conditions or classifications, such as calcium event categories, subnuclei locations, and clustered vs single profiles, etc. Both models included the same random effects to account for the hierarchical structure of the data, where observations were nested within higher-level units, such as neurons, recordings, etc.

The general form of the *full model* was:

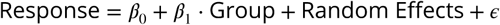

where *β*_0_ is the intercept, *β*_1_ is the coefficient for the group fixed effect, and the random effects account for variation within clusters (e.g., neurons and recordings). *ε* represents the residual error, capturing the variation in the response variable that is not explained by the fixed or random effects in the model.

The *reduced model* was constructed by excluding the group variable:

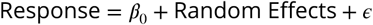

To determine if the group variable significantly contributed to the model, we performed a likelihood ratio test (LRT) to compare the fit of the full model to that of the reduced model by evaluating the difference in their log-likelihoods. A significant LRT would indicate that the group variable significantly influences the response variable. If the full model does not offer a better fit than the reduced model (i.e., the log-likelihood of the full model is not greater), it suggests that the group variable does not enhance the model fitting. In such cases, the LRT is not applicable, and consequently, a p-value is not provided, as the test relies on the assumption that the more complex model (full model) provides a statistically significant improvement in fit over the simpler model (reduced model).

Bootstrap test using 1000 itineraries was applied on peri-stimulatory calcium event frequency to evaluate the influence of stimulation on the temporal dynamics of neuronal activity.

To evaluate the clustering effect and probability of calcium spikes, we used the Wilcoxon signedrank test (signrank) or Mann-Whitney U test (ranksum) when the sample size is small. For all analyses, statistical significance was defined at levels (n.s.: *p* > 0.05, *: *p* < 0.05, **: *p* < 0.01, ***: *p* < 0.001).

## Supplementary video descriptions

**Video 1.** Calcium imaging video used in Figure 1 C; spontaneous activity of PO and DAO neurons shown as raw (tiff) images, denoised and deconvoluted versions.

**Video 2.** Calcium imaging video used in Figure 3 A; spontaneous and airpuff-ecoked activity in PO neurons shown as denoised and deconvoluted versions. Cyan frame represents the airpuff stimulation.

**Video 3.** Calcium imaging video used in Figure 4 B1 (PO); spontaneous activity suppression by N-O stimulation in PO neurons, shown as denoised and deconvoluted versions. Magenta frame represents the duration of the optogenetic stimulation.

**Video 4.** Calcium imaging video used in Figure 4 B1 (DAO); spontaneous activity suppression by N-O stimulation in DAO neurons, shown as denoised and deconvoluted versions. Magenta frame represents the duration of the optogenetic stimulation.

**Video 5.** Calcium imaging video used in Figure 5 A3 (PO); spontaneous activity suppression by N-O stimulation in DAO neurons and AP responses, shown as denoised and deconvoluted versions. Magenta frame represents the duration of the optogenetic stimulation; cyan frame indicates airpuff stimulation.

## Acknowledgments

We acknowledge the OIST Animal Resources Section (ARS) for support in animal well-being, and the Scientific Computing and Data Analysis (SCDA) section for computing facility. This research was funded by OIST intramural funding and KAKENHI grant 22K15207 to GD.

---

**Figure 2—figure supplement 1.**
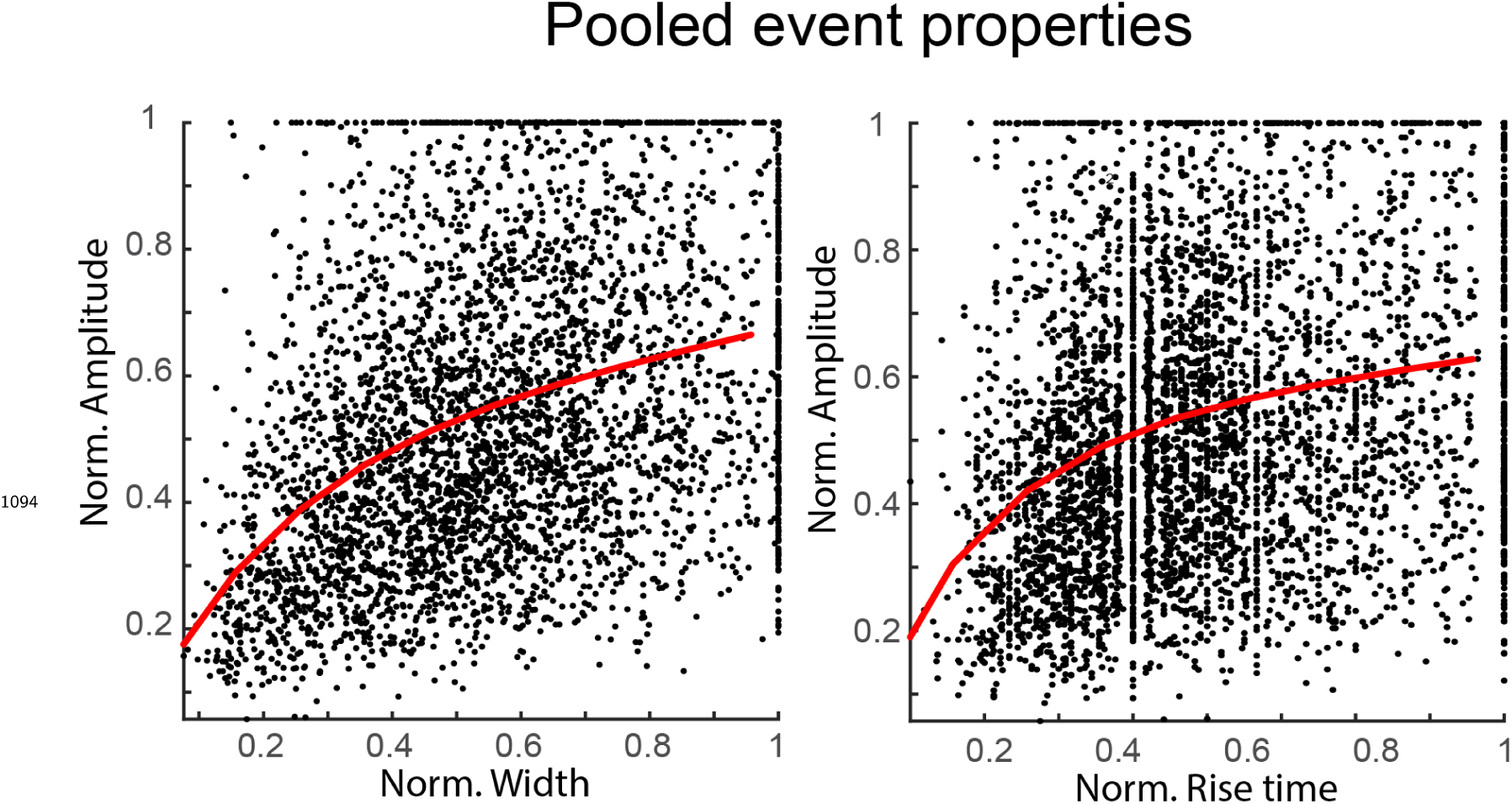
Comparison of event amplitudes and widths (left) and rise times (right), all shown normalized to the largest values within each cell. Amplitudes do not grow linearly throughout the range of event width, possibly due to GCamP6s saturation with largest events. However, with the low sampling frequency, amplitude is more reliable metric than duration.

**Figure 2—figure supplement 2.**
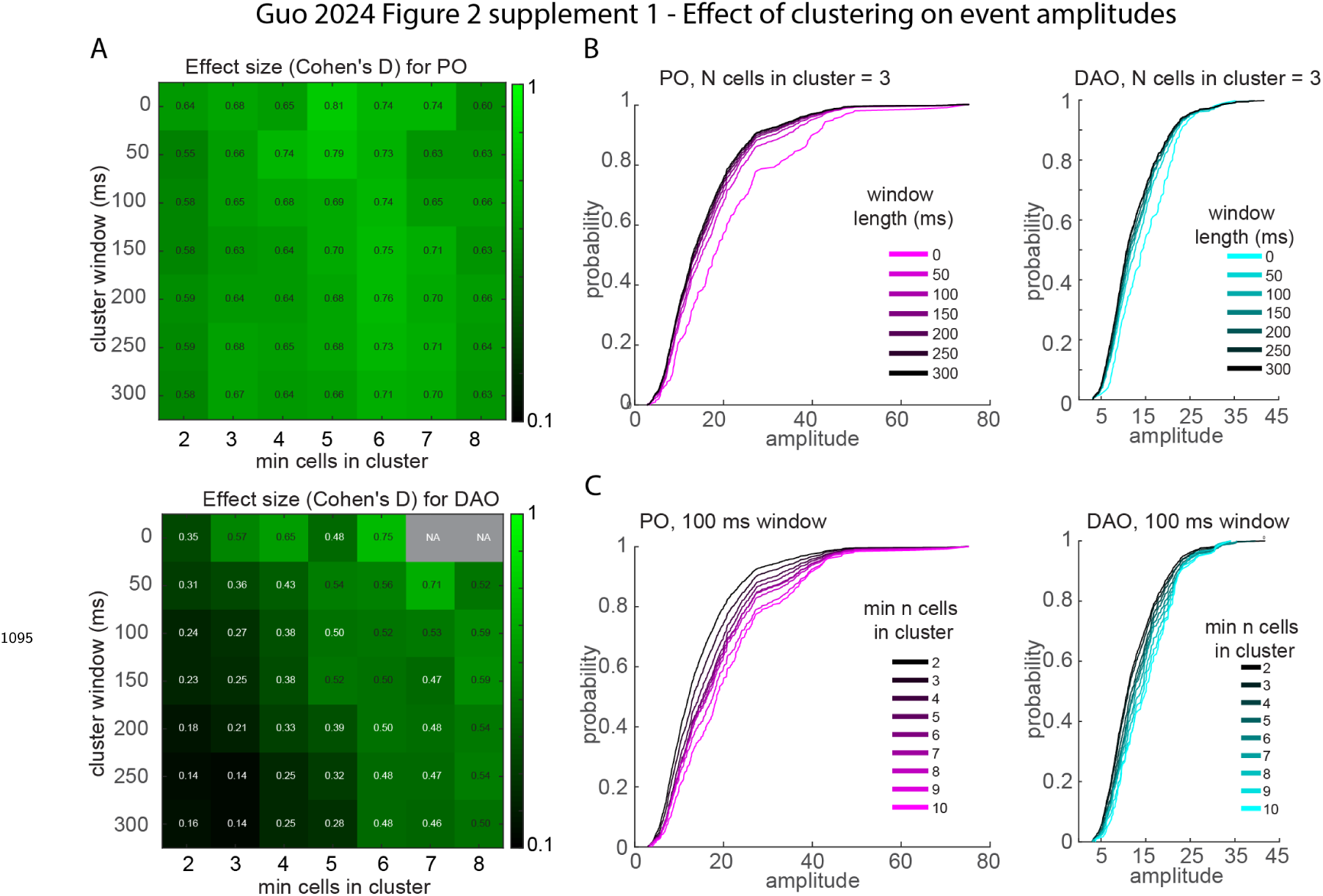
Parameter sweeps for clustering effect on event amplitude for PO and DAO. A, effect sizes on event amplitudes with clustering window (epsilon in DBSCAN algorithm) increasing from 0 300 ms (0-7 frames), and minimum number of events needed to form a cluster increasing from 2 to 7. Color brightness depicts effect size of the change in event amplitude distribution; effect sizes also indicated in squares. Gray squares indicate no available data. Highests effects are seen with short windows and moderate numbers of events in a cluster, as less data is available for large clusters. B, event amplitude distributions for 3-spike clusters are modulated by varying acceptable window lengths for clustering in PO (left) and DAO (right). C, event amplitude distributions for 100-ms windows are affected by varying cluster sizes in PO (left) and DAO (right).

**Figure 4—figure supplement 1.**
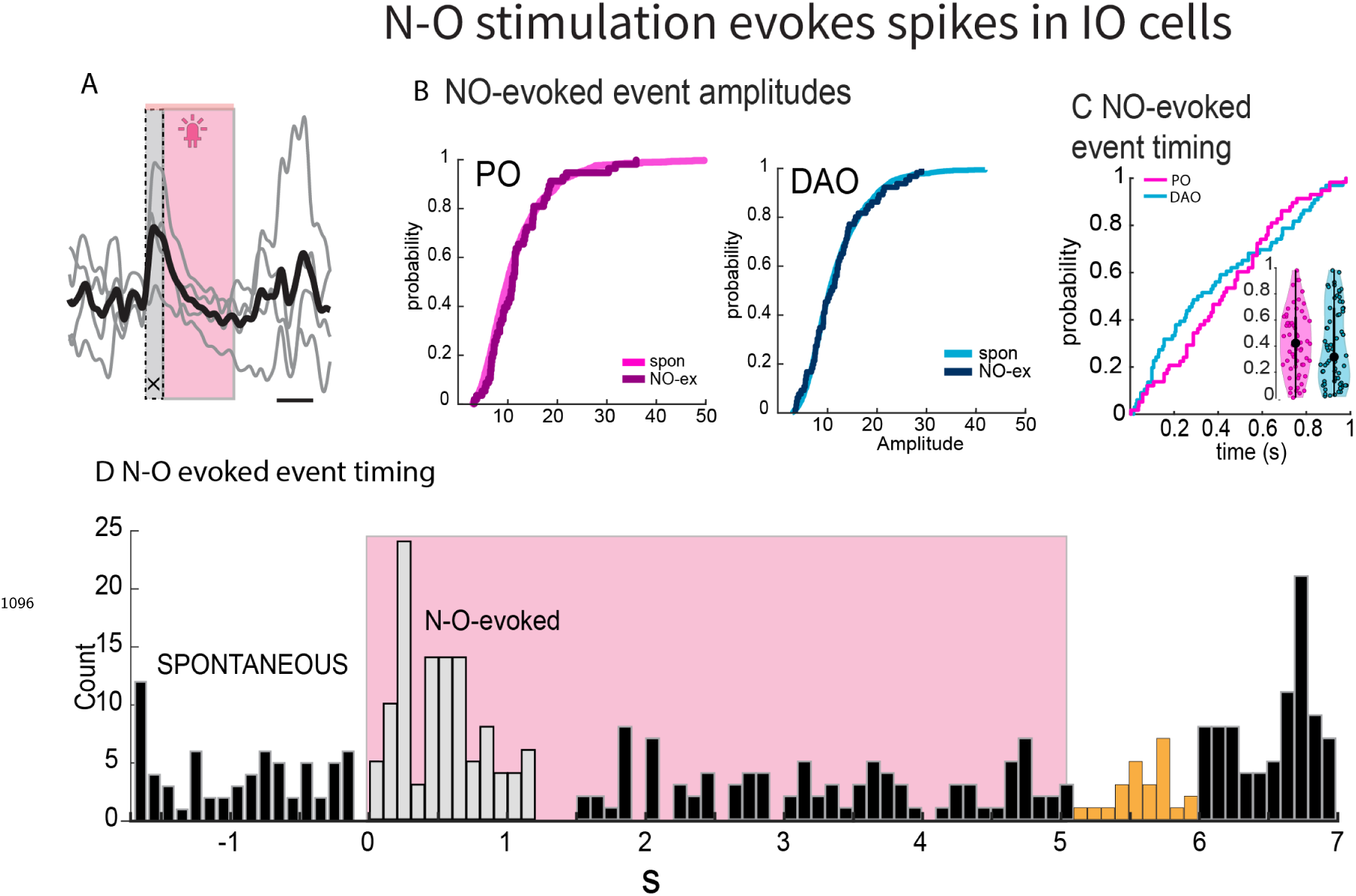
A, example traces from a single cell (4 stimulations) where optogenetic activation of N-O axons robustly leads to a spike at the onset of the stimulation. Gray box indicates the time window during which spikes were categorized as N-O-evoked. B, Cumulative histograms of NO-evoked spike amplitudes vs. the spontaneous events in the same trials in PO and DAO. C, timing of N-O-evoked events with respect to onset of optogenetic stimulation. D event frequency histogram time-aligned at the onset of N-O stimulation from trials where a N-O evoked spike was seen (white bars).

**Figure 4—figure supplement 2.**
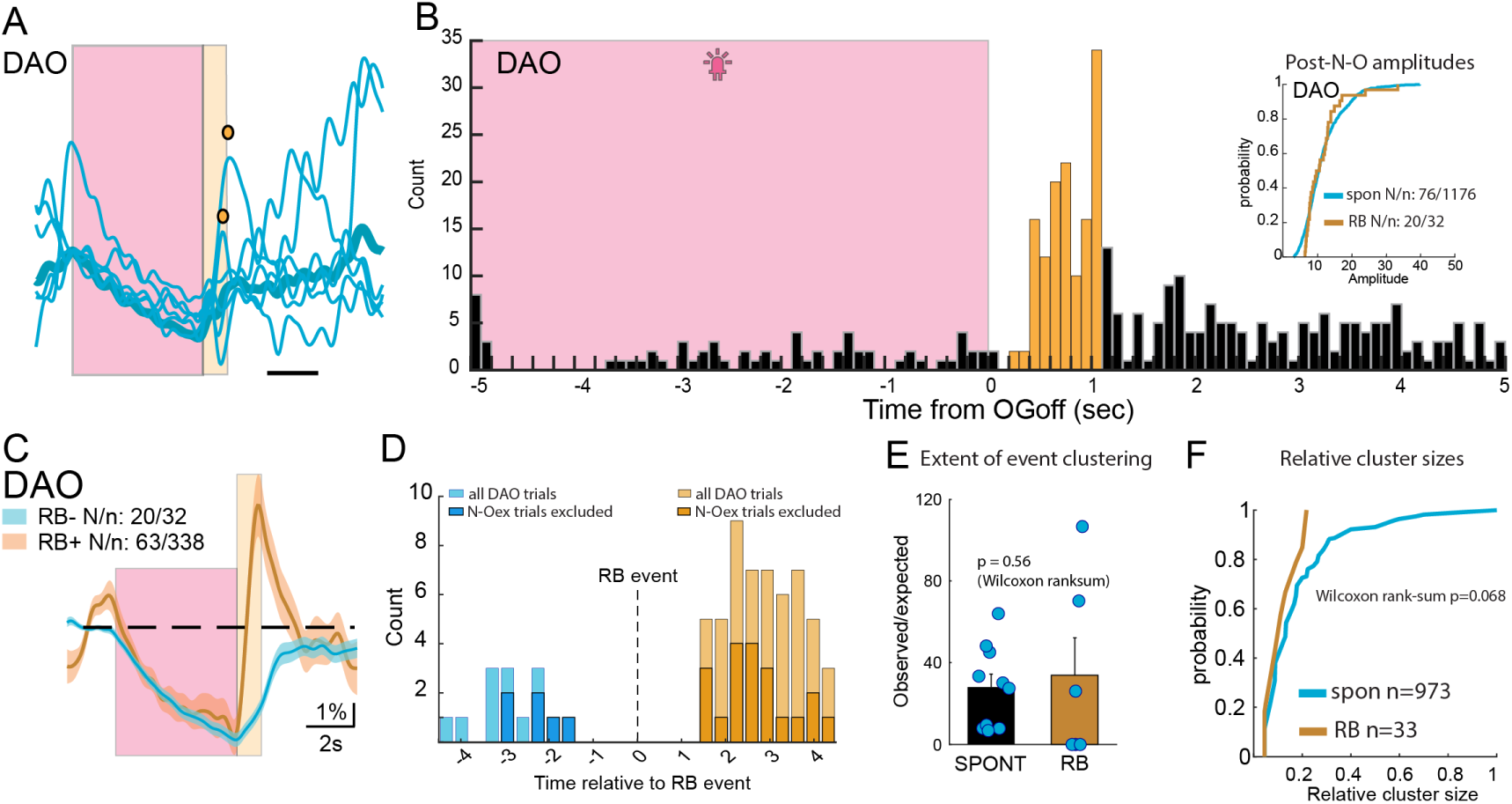
A, examples of DAO cell stimulation trials where “rebound events” (RB) were seen in the post-stimulation window (indicated in yellow color). Circles denote detected events. B, histogram of the frequency of the event during N-O stimulation and 5 seconds following it in trials with RB events (yellow bars). The inset shows comparison of the amplitudes of RB events to spontaneous events. C, mean calcium traces from N-O stimulation trials in all DAO cells, grouped by whether an event occurred within the rebound window (yellow) or not (blue), Thich trace depicts average trace, shading±SSEM. No difference in the amplitude of calcium suppression during N-O stimulation is seen between these groups. D, timing of events preceding (blue) and following (brown) the RB event. Due to low number of events, data from trials where N-O stimulation led to an early spike are included (light-colored bars); data from trials after N-O driven spikes are excluded are shown in dark colors. E, F: RB events do not show difference with respect to event clustering probability (E) or cluster sizes (F). E3, comparison of synchronous and asynchronous amplitudes for post-stimulation events. E4, distribution of delays of post-stim spikes with respect to the offset of optogenetic stimulus (“OGoff”), compared to the distribution of inter-event intervals among spontaneous events.

## Notes

### Competing Interest Statement

The authors have declared no competing interest.

